# Identification of echinacoside as a tobramycin potentiator against *Pseudomonas aeruginosa* aggregates

**DOI:** 10.1101/2024.02.09.579617

**Authors:** Yu-Ming Cai, Aurélie Crabbé, Tom Coenye

## Abstract

Cyclic diguanylate (c-di-GMP) is a central biofilm regulator, where increased intracellular levels promote biofilm formation and antibiotic tolerance. Targeting the c-di-GMP network is a promising anti-biofilm approach. Most agents reported previously decreased c-di-GMP to eliminate surface-attached biofilms, which did not recapitulate *in vivo* biofilms well and may thus impede their clinical impact. Here, the expression profile of genes encoding proteins associated with c-di-GMP metabolism was analysed among 32 *Pseudomonas aeruginosa* strains grown as suspended aggregates in synthetic sputum or planktonic cells. A diguanylate cyclase, SiaD, proved essential for auto-aggregation under *in vivo*-like conditions. Virtual screening against SiaD identified echinacoside as an inhibitor, which reduced intracellular c-di-GMP levels and aggregate sizes and potentiated the efficacy of tobramycin against aggregates established by >80% of tested strains. This synergistic effect was also observed for *in vivo*-like 3-D alveolar cells infected by cytotoxic *P. aeruginosa*, demonstrating its high potential as an adjunctive therapy for recalcitrant *P. aeruginosa* infections.

## Introduction

It is estimated that biofilms, surface-attached or suspended multicellular bacterial communities embedded in a self-produced matrix, are associated with more than 80% of chronic infections^1,2^. The secretion of extracellular polymers (including polysaccharides, proteins, and DNA), reduced metabolic activity, and an altered microenvironment all contribute to the reduced susceptibility of biofilm-associated microorganisms towards antimicrobial agents and host responses^3,4^ and render the eradication of chronic infections difficult at best^5^. With the slow progress in antibiotic discovery, novel anti-biofilm strategies that enhance the susceptibility of microbial cells towards conventional antibiotics are urgently needed^6^.

Different therapeutic approaches are under development to either interfere with crucial structural traits of biofilms or modulate signalling pathways that regulate biofilm formation and dispersal^7,8^. The intracellular secondary messenger c-di-GMP emerged as a promising target due to its central roles in biofilm regulation and wide distribution among bacterial species^9,10^. An increased production of c-di-GMP promotes biofilm formation and maturation through multiple mechanisms, and conversely a reduction in intracellular c-di-GMP levels inhibits biofilm or triggers dispersal^9^. C-di-GMP is synthesized by diguanylate cyclases (DGCs) with a core GG(D/E)EF catalytic domain and is hydrolyzed by phosphodiesterases (PDEs) with either E(A/V)L or HD-GYP core domains^10^. As such, inhibition of DGCs or stimulation of PDEs is considered a promising way to decrease c-di-GMP levels and reduce biofilms^11^.

However, the c-di-GMP signaling pathway is a highly complex network, where multiple genes encoding DGCs and PDEs collectively contribute to modulating the c-di-GMP concentration in response to a myriad of environmental stimuli^12,13^. Not all of these proteins influence c-di-GMP levels in the same way: some DGCs and PDEs alter the overall level, while others fine-tune local concentrations^13,14^. In addition, the production or hydrolysis of c-di-GMP may occur in a stepwise manner throughout the course of biofilm development, where different DGCs/PDEs function as the predominant regulators in each distinct stage^15^. The plasticity and sophistication of this highly organized pathway hence present a major challenge for the development of effective small molecules that interfere with c-di-GMP signalling.

A number of compounds that inhibit or disperse surface-associated biofilms by inhibiting DGCs, stimulating PDEs, controlling riboswitches, and sequestering c-di-GMP by direct binding have been identified^8,16–18^. However, while such surface-associated biofilms have been extensively studied, more recent evidence demonstrated that biofilms in chronic infections predominantly form suspended (non-attached) aggregates surrounded by host secretions and inflammatory cells^19–21^. While surface-attached biofilms and non-attached aggregates share similar phenotypes^19^, it is currently unclear whether agents targeting surface-attached biofilms can also inhibit non-attached aggregates in more complex *in vivo*-like conditions. To this end, we determined to discover drugs targeting c-di-GMP, and more specifically, novel DGC inhibitors, to reduce auto-aggregation of *P. aeruginosa*, a widespread opportunistic human pathogen associated with many diseases, of which the most well-documented is chronic lung infections in cystic fibrosis (CF) patients. To generate a more *in vivo-*like environment, *P. aeruginosa* aggregates were grown in synthetic cystic fibrosis medium 2 (SCFM2), which is regarded as the most accurate pre-clinical model developed to mimic CF mucus so far^22^. In *P. aeruginosa*, 42 proteins were reported to contain GG(D/E)EF, EAL/HD-GYP, or dual domains^23,24^, of which WspR has been used as the target for all previously identified DGC inhibitors in *P. aeruginosa*. However, we found that deleting *wspR* in PAO1 did not affect the overall auto-aggregation level in SCFM2. Therefore, a comprehensive RT-qPCR assay was performed to compare the gene expression profile of 40 genes encoding DGCs and PDEs between aggregated and planktonic cells of 32 *P. aeruginosa* clinical isolates. Such a large-scale assay and further gene function analysis led to the identification of SiaD, a previously reported DGC responsible for *P. aeruginosa* auto-aggregation under the challenge of detergent^25,26^, as a key player also in aggregation in patient-mimetic conditions. By *in sillico* screening of 21495 bioactive compounds against SiaD, a natural phenylethanoid glycoside, echinacoside, was selected and shown to reduce global c-di-GMP level, inhibit aggregation in SCFM2, and increase the efficacy of tobramycin against pre-established aggregates formed by 27 out of 32 isolates by 2.7 to 48-fold. Furthermore, a synergistic effect of echinacoside and tobramycin was also observed for treating PA14 aggregates attached to 3-D human A549 cells that recapitulate *in vivo* tissue^27–29^. As such, our study not only identified an essential DGC responsible for *P. aeruginosa* auto-aggregation in CF sputum environment suitable for drug development, but also discovered a compound with high potential for improving *P. aeruginosa* treatment outcome in future clinical applications.

## Results

### SiaD (PA0169) is identified as an essential DGC responsible for auto-aggregation in SCFM2

WspR is a well-characterized active DGC associated with c-di-GMP production, surface sensing and biofilm formation, motility, as well as cell envelope stress^30–33^. However, its role in the formation of non-surface attached *P. aeruginosa* aggregates in more clinically relevant models is unclear. We observed that the lack of WspR did not impact the auto-aggregation of PAO1 in SCFM2 (Fig. 1a). The sizes of aggregates were comparable between WT and *ΔwspR*, where aggregates with sizes between 1000 μm^3^ and 50000 μm^3^ constituted 68.72±15.54% and 71.02±9.73% of total biovolume in WT and *ΔwspR* cultures respectively (Fig. 1b and c, Table S1). Subsequently, we evaluated the expression levels of 40 genes encoding enzymes involved in c-di-GMP metabolism in 32 *P. aeruginosa* clinical strains (Table S2). Genes included encode DGCs, PDEs and DGC-PDE dual domain proteins. *arr* and *pvrR* were excluded due to their absence in multiple strains (Table S3). We compared gene expression levels between aggregates grown in SCFM2 and planktonic cells grown in SCFM2 without DNA and mucin as planktonic cells, as mucin and DNA supports auto-aggregation more as physical rather than nutritional factors^34–36^ (Fig. S1a-c). As the growth phase significantly influences the expression patterns of c-di-GMP-related genes^37^, we cultivated different strains for different periods of time in order for them to reach the same growth phase (i.e. late log-phase) (Table S4). The RT-qPCR results did not identify any gene of which the expression fold change (aggregate *versus* planktonic cells) was substantially higher or lower than other genes among the majority of strains, where the fold-change values of most genes ranged from ∼0.2 to ∼2 (with a few exceptions in strains DK2, NH57388A, OS4, and BS6) (Fig 1d, supplemental RT-qPCR dataset). This finding was consistent with results from our ELISA-based intracellular c-di-GMP quantification assay (Fig. S2), where only 9 out of 32 strains showed a lower intracellular c-di-GMP level in planktonic cells compared to their aggregated counterparts, and the decrease was not as substantial as reported previously^38,39^. Despite the discrepancy between results and our initial hypothesis, *proE*, *fimX, PA4396*, *PA2567*, *siaD* and *PA5442* were ranked higher than others in the heatmap, indicating that the overall expression fold changes (aggregates *versus* planktonic cells) of these genes among different strains were higher than those of other genes. PA4396 is a degenerated DGC^40^; PA2567 is an EAL-containing PDE^41^; ProE harbours a degenerate GGDEF domain and exhibits PDE activity^42^; and FimX contains GGDEF and EAL domains that are both inactive^43^. SiaD was shown to be an active DGC involved in *P. aeruginosa* auto-aggregation in response to environmental stress^25,26,44^ and PA5442 was predicted to contain an active DGC domain^45^. To assess the role of these two proteins in auto-aggregation in SCFM2, isogenic mutants *ΔsiaD* and *ΔPA5442* were constructed in PAO1. Deleting *PA5442* did not significantly affect the aggregation level, as 71.06±10.37% and 68.54±8.45% of total biovolume were constituted by aggregates with sizes ranging from 1000 μm^3^ to 50000 μm^3^ in WT and *ΔPA5442* cultures, respectively. On the other hand, the lack of SiaD substantially reduced the size of aggregates, with aggregates larger than 1000 μm^3^ only making up 3.31±6.42% of the total biovolume (Fig. 1e-g, Table S1). Complementing *ΔsiaD* with *siaD* restored the aggregative behaviour (Fig. S3 and Table S1). Therefore, we concluded that *siaD* is essential for aggregate formation in SCFM2 and SiaD was selected for further studies.

**Fig. 1.**
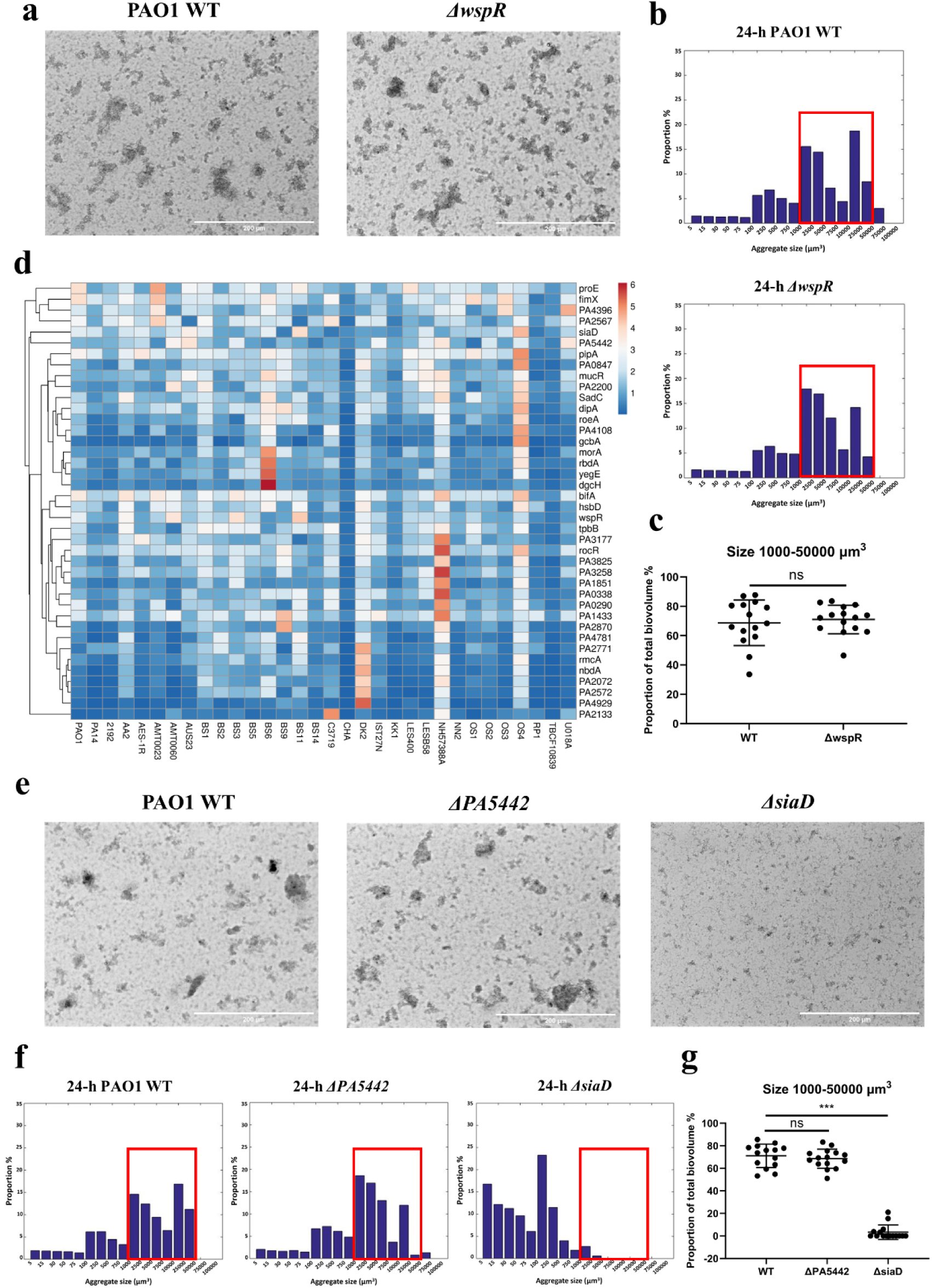
**(a)** Representative micrographs of *P. aeruginosa* PAO1 WT and *ΔwspR* aggregates grown for 24 hrs in SCFM2. Scale bar = 200 µm. **(b)** Distribution of the size of 24-h PAO1 WT and *ΔwspR* aggregates grown in SCFM2. The biovolume (µm^3^) of single cells and aggregates were grouped into 17 categories. The biovolume of all aggregates belonging to each size category was calculated, and the proportion of this biovolume of each category in the total biovolume was calculated. Histograms represent the mean proportion value of all samples collected from 3 individual experiments (n=3), where micrographs were obtained from at least 3 random locations in each sample. Numerical data and standard deviations are shown in Table S1. **(c)** Proportion of aggregates with sizes ranging from 1000 to 50000 μm^3^ in total biovolume. ***, p<0.001 (Student’s t-test) **(d)** Heat map showing the relative expression level (fold changes) of genes within the *P. aeruginosa* c-di-GMP network (*y*-axis) in aggregated cells *versus* planktonic cells of 32 strains (*x*-axis). Unit variance scaling is applied to rows. Imputation is used for missing value estimation. Rows are clustered using correlation distance and average linkage. Tree for rows were ordered where genes with higher mean values were shown first. **(e)** Representative micrographs of PAO1 WT, *Δpa5442*, and *ΔsiaD* aggregates grown for 24 hrs in SCFM2. Scale bar = 200 µm. **(f)** Distribution of the size of 24-h PAO1 WT, *Δpa5442* and *ΔsiaD* aggregates grown in SCFM2. Histograms represent the mean proportion value of all samples collected from 3 individual experiments (n=3), where micrographs were obtained from at least 3 random locations in each sample. Numerical data of mean proportions and standard deviation are shown in Table S1. **(g)** Proportion of aggregates with sizes ranging from 1000 to 50000 μm^3^ in total biovolume. ***, p<0.001; ns, not significant (Student’s t-test).

### Echinacoside reduces both auto-aggregation and intracellular c-di-GMP levels of PAO1 in SCFM2

The structure of SiaD deposited in PDB (code 7E6G) was used as the template for virtually docking 21495 compounds with known bioactivities into the conserved GTP substrate-binding site (GGDEF domain, A-site). The docking score refers to the binding affinity between these compounds and the coordination pocket of SiaD, with lower docking scores correlating with higher binding affinity and generally a score lower than −10 (−10 kcal/mol) suggests a highly promising binding. Results showed that bimosiamose and echinacoside exhibited the highest binding affinity among 21495 compounds, with docking scores of −10.260 and −10.259 respectively (Fig. 2a, Supplemental Excel top 200 compounds identified by virtual docking). Bimosiamose can form six hydrogen bonds with ASP103, ARG177, GLY180, GLU182, and VAL139, and the benzene ring can form one cation-π interaction with LYS143. Multiple hydroxyl groups of echinacoside can form nine hydrogen bonds with ASP155, GLU182, GLY180, ASP243, VAL139, LYS143, and ARG254 (Fig. 2a, upper panel). While neither of these two compounds showed a bactericidal effect at tested concentrations (Fig. S4), only echinacoside successfully reduced the aggregation level of PAO1 in SCFM2. When PAO1 was incubated with 250 nM echinacoside, only 11.28±9.68% of bacterial cells grew into aggregates larger than 1000 μm^3^. In contrast, 43.61±7.94% of the total biovolume of untreated cells was composed of aggregates larger than 1000 μm^3^ (Fig. 2b-d, Table S1). The effect of echinacoside on aggregation of PAO1 in SCFM2 was concentration-dependent, and the reduction was abolished when echinacoside was applied at concentrations higher than 1 µM with the initial inoculum size being 1×10^5^ CFU/mL (Fig. S5 and Table S1). While the mechanism underlying this phenomenon requires further investigations, we applied echinacoside at a final concentration of 50 µM to 1×10^7^ CFU/mL PAO1 and *ΔsiaD* cells to evaluate if echinacoside can reduce intracellular c-di-GMP level. As shown in Fig. 2e, echinacoside reduced the c-di-GMP level in PAO1 WT but not in *ΔsiaD* cultured in SCFM2, suggesting the interaction between SiaD and echinacoside.

**Fig. 2.**
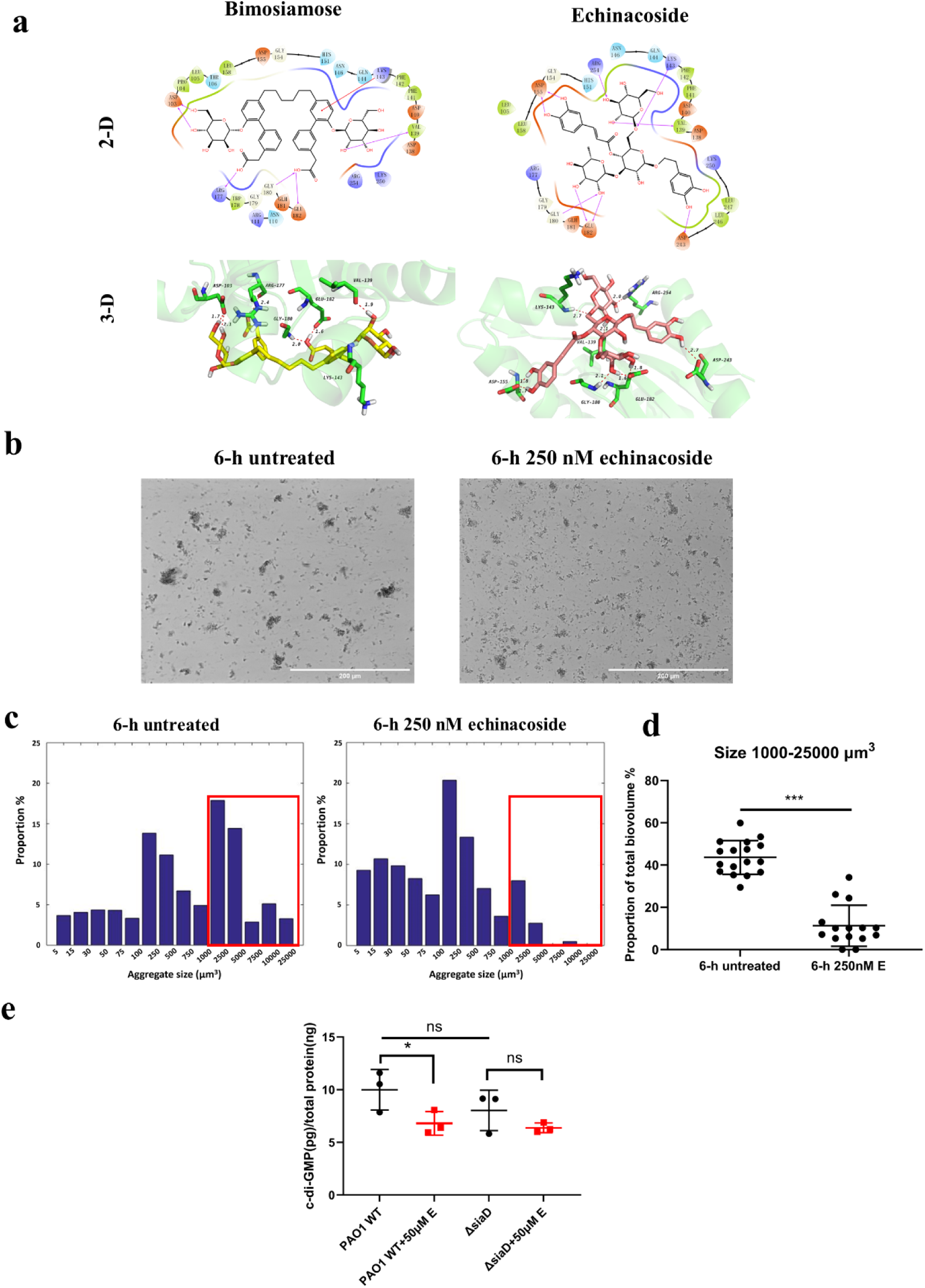
**(a)** Interaction between two compounds with the lowest *in silico* docking score and SiaD. Chains B/C/D/E/F, water molecules, magnesium ions and the missing hydrogen atoms of SiaD were deleted. In the 3-D illustration (lower panel), the carbon skeleton of SiaD is displayed in green, N atoms in blue, O atoms in red, H atoms in white, and hydrogen bonds as red dashed lines. In corresponding images, bimosiamose and echinacoside were displayed as light yellow and dark pink sticks, respectively. The distances of hydrogen bonds are also shown. **(b)** Representative micrographs of *P. aeruginosa* PAO1 aggregates grown for 6 hrs in SCFM2 with or without 250 nM echinacoside. Scale bar = 200 µm. **(c)** Distribution of the size of 6-h PAO1 WT aggregates grown in SCFM2 with or without 250 nM echinacoside. The biovolume (µm^3^) of single cells and aggregates were grouped into 14 categories. Histograms represent the mean proportion value of all samples collected from 3 individual experiments (n=3), where micrographs were obtained from at least 3 random locations in each sample. Numerical data of mean proportions and standard deviation are shown in Table S1. **(d)** Proportion of aggregates with sizes ranging from 1000 to 25000 μm^3^ in total biovolume. ***, p<0.001 (Student’s t-test). **(e)** Intracellular c-di-GMP levels in *P. aeruginosa* PAO1 WT and *ΔsiaD* grown in SCFM2 with or without 50 µM echinacoside treatment normalized to total protein concentration. Mean values of 3 independent experiments with 2 technical replicates each; error bar denotes standard deviation. *, p<0.05; ns, not significant (Student’s t-test).

### Echinacoside potentiates the activity of tobramycin against *P. aeruginosa* aggregates formed by clinical isolates

As echinacoside successfully inhibited the formation of aggregates, we next assessed whether it could potentiate the activity of frequently used antibiotics. Echinacoside was first applied to pre-established PAO1 aggregates in SCFM2 along with 4 antibiotics belonging to different categories (tobramycin, ciprofloxacin, meropenem and ceftazidime). Echinacoside enhanced the antimicrobial activity of both tobramycin and meropenem, but not the activity of ciprofloxacin or ceftazidime (Fig. 3a and b, Fig. S6). When this screening was expanded to 7 additional *P. aeruginosa* strains, the potentiating effect towards tobramycin was more consistent than towards meropenem (for certain strains, the addition of echinacoside even appeared to weaken the bactericidal effect of meropenem; Fig. S7) and we selected tobramycin for further experiments. Different combinations of echinacoside (25, 50, 100, 200, 400, 800 µM) and tobramycin (at least 2 different concentrations leading to ≥3-log reduction in CFU) were tested on pre-established aggregates of 32 *P. aeruginosa* strains grown for different periods in SCFM2 (Table S4) to optimize concentration ranges. For 27/32 strains (BS3, BS9, BS11, OS4 and C3719 being the exceptions), the addition of echinacoside led to an extra 2.7 to 48-fold reduction compared to treatment with tobramycin alone (Fig. 3c, Fig. S8). It is currently unclear why echinacoside did not potentiate tobramycin activity against strains BS3, BS9, BS11, OS4 and C3719. BS3, BS9, and C3719 are resistant to tobramycin (Table S5; EUCAST breakpoint for resistance is 2 µg/mL), but echinacoside did potentiate tobramycin activity against other tobramycin resistant strains (e.g. BS5 and DK2). Likewise, BS11 and OS4 are highly mucoid but echinacoside did potentiate tobramycin activity against other mucoid strains (e.g. BS1 and BS5).

**Fig. 3.**
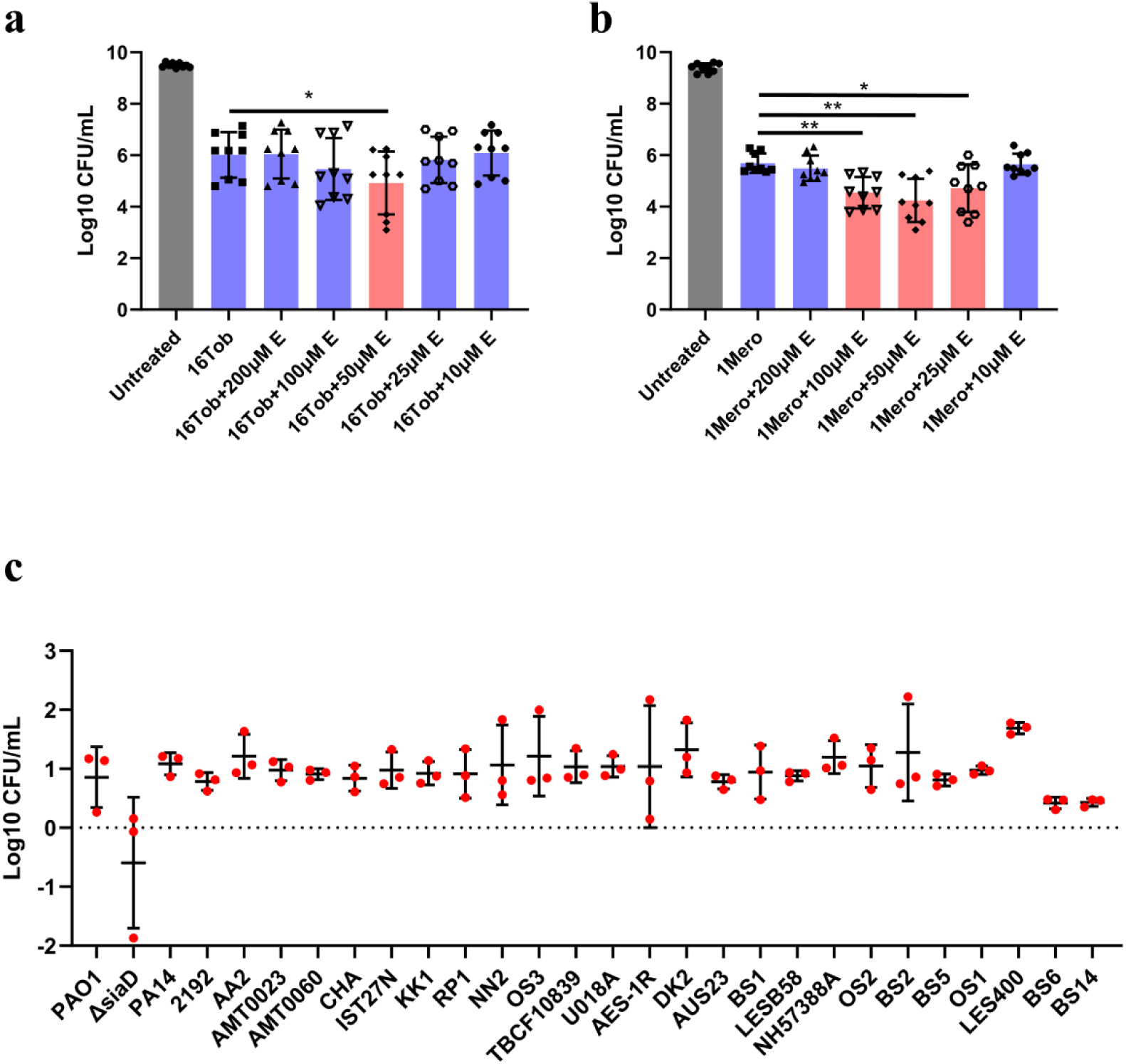
**(a,b)** The efficacy of 16 μg/mL tobramycin (16Tob) **(a)** and 1 μg/mL meropenem (1Mero) **(b)** in combination with different concentrations of echinacoside (10 μM to 200 μM E) against pre-established (6-h) PAO1 aggregates grown in SCFM2. Data are expressed as the mean number of CFU remaining after an additional 18-h incubation (3 independent experiments with 3 technical replicates; error bars indicate standard deviation). Red bars highlighted the successful combination treatment that potentiated the efficacy of antibiotics. *, p<0.05; **, p<0.01 (Student’s t-test). **(c)** Log reduction achieved by combination treatment of pre-established aggregates of 28 *P. aeruginosa* strains, compared to treatment with tobramycin alone was shown. (3 independent experiments with 3 technical replicates; error bars indicate standard deviation).

### Echinacoside potentiates the efficacy of tobramycin in a physiologically relevant 3-D model of human alveolar epithelial cells

We subsequently evaluated if the synergistic effect could also be observed in a 3-D model of human alveolar epithelial (A549) cells that recapitulated some key aspects of *in vivo* tissue^27,29^. The widely used reference strain PA14 was chosen for infection assays due to its higher virulence compared to PAO1 and well-documented cytotoxicity against A549 cells^46–49^. After 4 hr co-culture of 3-D A549 cells and PA14, aggregates were found attached to the apical cell surface (Fig. S9a). Echinacoside was not cytotoxic to 3-D A549 cells at concentrations ranging from 25 µM to 400 µM based on both LDH assay results and micrographs shown in Fig. S9b and c. We then assessed the viability of PA14-infected 3-D A549 cells with different treatments. While the survival of PA14-infected cells with echinacoside or 2 μg/mL tobramycin treatment alone were comparable to those without treatment, cells treated with 25 or 50 µM echinacoside together with tobramycin showed significantly higher survival compared to untreated cells or those treated by tobramycin alone (Fig. 4a). Such outcome can also be observed microscopically, where healthy cells firmly attached to collagen-treated microcarrier beads exhibited a smooth morphology and stressed or dead cells detached from beads form cell aggregates or floated in the medium. As shown in Fig. 4b, when infected A549 cells were treated with both tobramycin and echinacoside, more cells remained attached to beads compared to cells treated with tobramycin alone. In conclusion, micromolar scale echinacoside was not cytotoxic, and a synergistic effect of echinacoside and tobramycin was observed against *P. aeruginosa* aggregates formed on 3-D A549 cells.

**Fig. 4.**
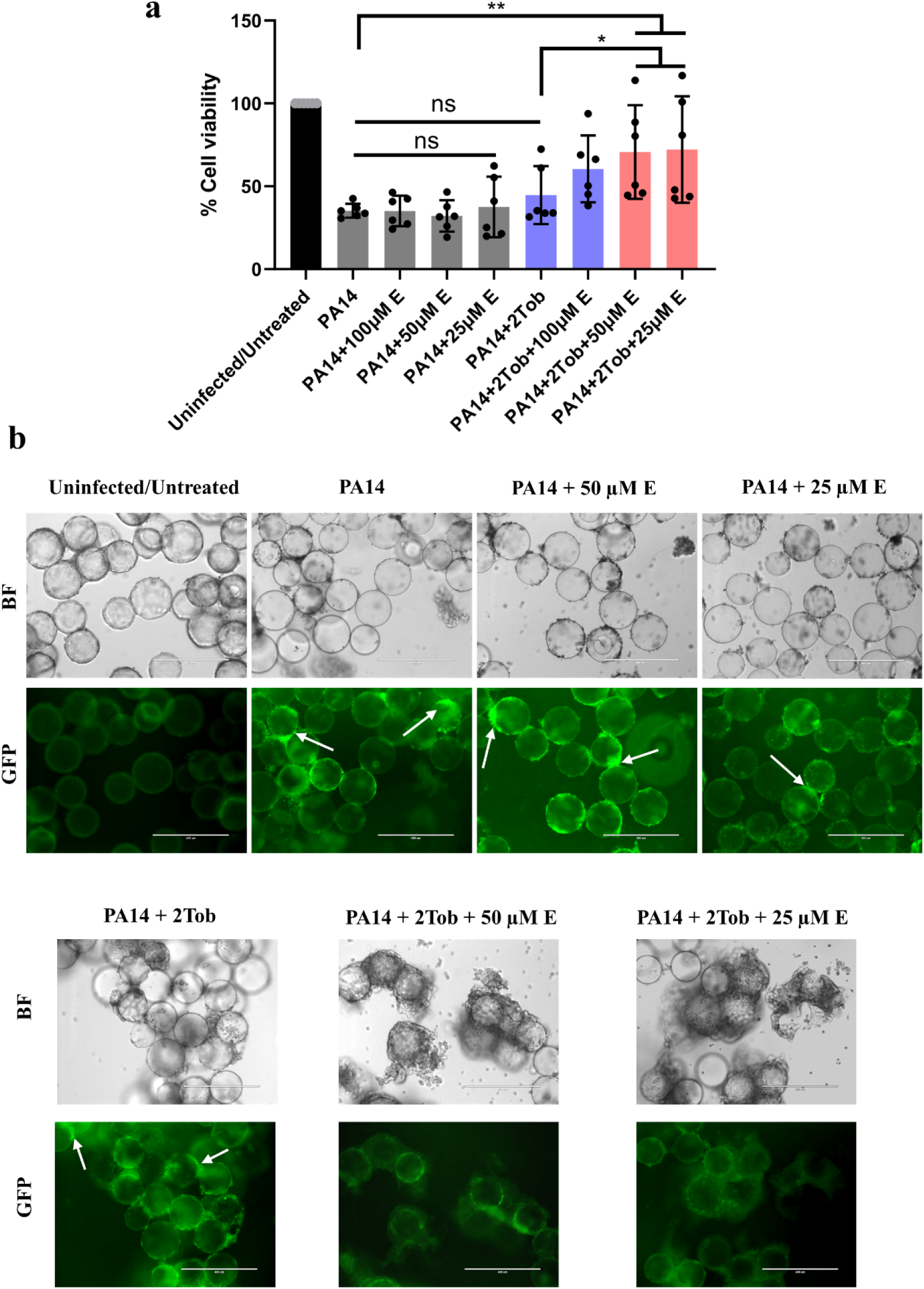
**(a)** Viability of PA14-infected 3-D A549 cells (as measured with an LDH assay) exposed to different concentrations (25, 50 and 100 μM) of echinacoside, 2 μg/mL tobramycin (2Tob) or the echinacoside/tobramycin combination treatments normalized to untreated/uninfected groups. Red bars highlighted the successful combination treatment that potentiated the efficacy of tobramycin and elevated cell viability. (6 independent experiments with 4 technical replicates; error bars indicate standard deviation). *, p<0.05; **, p<0.01 (Student’s t-test). **(b)** Representative transmission light micrographs (upper panel) of 3-D A549 cells attached to/detached from collagen-treated carrier beads and fluorescent micrographs (lower panel) of GFP-tagged PA14 attached to A549 cells. Some typical PA14 aggregates attached to A549 cells with strong fluorescent signals are highlighted with white arrows. Scale bar = 400 μm.

## Discussion

In the present study, we showed that WspR, the most-studied *P. aeruginosa* DGC so far, does not play a role in determining aggregate size in SCFM2 (Fig. 1a, b, c). This is not surprising, as the Wsp signal transduction complex is a chemosensory system responding to the growth of *P. aeruginosa* on surfaces^30,50^. Multiple studies have shown that constitutive activation of WspR through loss/mutation of WspF or overexpression of *wspR* significantly increases c-di-GMP levels and biofilm formation.^30,51–53^. However, despite its thoroughly proven DGC activity^30,54^, lacking WspR *per se* did not influence the overall biomass but more on biofilm 3-D structure^23,24^. Therefore, the inactivation of WspR in strains without mutations in *wspF* may not be the optimal strategy solely from the perspective of its role in biofilm formation. Mutations in *wspF* were claimed to be responsible for the emergence of frequently isolated *P. aeruginosa* rugose small-colony variants (RSCVs) from chronic infections, which lead to hyper-biofilm formation and persistent phenotypes^55,56^. However, the frequency of *wspF* mutation is relatively low compared to some mutation hotspots in clinical isolates adapted in patients, and more interestingly, lower than *wspA* and *wspE* in the same family^57–59^. Besides, another operon, *yfiBNR*, also contributes to RSCV development^55,60^. Combining these previous findings and our results, it is reasonable to conclude that we need to search for other c-di-GMP catalysts as targets to reduce aggregation *in vivo*.

We subsequently quantified the expression of 40 genes in the c-di-GMP network in aggregated and planktonic *P. aeruginosa* cells, expecting to identify one or a few DGC-encoding genes expressed at a significantly higher level than others for discovering DGC inhibitors. Due to both the genetic and physiological versatility of *P. aeruginosa*, 32 strains, including 2 reference strains PAO1 and PA14 as wound isolates and 30 CF isolates from different sources (Table S2), were used to better summarize the general expression pattern among the community. Specifically, aggregates were grown under microaerophilic environment providing oxygen level similar to those experienced by some *P. aeruginosa* strains *in vivo* and contributing to increased antibiotic tolerance^61^. However, our large-scale RT-qPCR analysis did not select an outstanding DGC or PDE with an expression pattern much different from others (Fig. 1d), which is consistent with the data where only 28% tested strains showed a decreased c-di-GMP level in planktonic cells compared to aggregates (Fig. S2). Such discrepancy between our data and the previous general belief that the global c-di-GMP levels in mature *P. aeruginosa* biofilms are at least 2 to 3-fold higher than planktonic counterparts^38,39,62–64^ may be explained by different nutrition backgrounds (LB or minimal medium in other studies), different culturing time and sampling techniques where we inevitably included planktonic cells for aggregate samples, as well as different metabolic rates in oxygen-limited environments. Due to the different intrinsic catalytic activity of each enzyme and the fact that c-di-GMP can be modulated both transcriptionally and post-translationally^65^, we propose that even a small increase in the gene expression level across different aggregated samples may suggest an interesting target. As such, we tested whether the loss of *PA5442* and *siaD*, two genes predicted or proved to encode DGCs respectively^45,66^, can reduce aggregation. Results indicated that SiaD does play an important role in auto-aggregation in SCFM2 (Fig. 1e-g), which corresponds to its contribution to auto-aggregation in other liquid environments^25,67^, and thus served as our drug docking template in the search for novel DGC inhibitors.

Both promising compounds identified through virtual screening contain multiple hydroxyl groups (Fig. 2a), a property sometimes disadvantageous for drugs, which is also consistent with many previously identified DGC inhibitors^16^. However, echinacoside was repeatedly reported as a potential neuroprotective, hepatoprotective and anti-cancer agent in animal models^68^, and in our case successfully inhibited auto-aggregation in such a complex medium difficult for molecule diffusion^69^ (Fig. 2b-d), indicating its high potential for clinical applications. Though deleting *siaD* did not lower the intracellular c-di-GMP level to a statistically significant degree as previously shown^44,67^ probably due to similar reasons discussed before, a slight decrease was observed, and echinacoside successfully reduced c-di-GMP level in PAO1 WT but not in *ΔsiaD* mutant (Fig. 2e). Despite the lack of data from kinetic analysis showing whether echinacoside interferes with the catalytic activity of SiaD exclusively or can inhibit multiple DGCs, our results are suggestive that echinacoside may regulate global c-di-GMP *via* SiaD.

Of course, one of the most important goals of developing anti-biofilm strategies is re-sensitizing biofilm-embedded cells towards conventional antibiotics. Echinacoside was recently demonstrated to inhibit biofilm formation and potentiate vancomycin against methicillin-resistant *Staphylococcus aureus* by binding to a transpeptidase^70^. Here, we first demonstrated that a synergistic killing effect of echinacoside and tobramycin can be observed for aggregates established by >80% tested CF isolates (including resistant and mucoid strains) in mucus-like environments, with sizes similar to most of those observed in CF patients (5-100 μm in diameter)^21^ (Fig. 3c, Fig. S8 and Table S5). Notably, this potentiating effect was not observed on *ΔsiaD* in SCFM2, which is consistent with c-di-GMP measurement results and further hints at the interference of echinacoside on SiaD activity (Fig. 3c and S8). Moreover, echinacoside also potentiated the efficacy of tobramycin against *P. aeruginosa* biofilms on the surface of 3-D A549 cells and protected against *P. aeruginosa*-induced cytotoxicity (Fig. 4). These positive data suggested that echinacoside has a high potential as an adjunctive therapy to treat *P. aeruginosa* embedded in the CF airway mucus layer. Intriguingly, for aggregates grown in both SCFM2 and 3-D A549 cell models, this potentiating effect was heavily concentration-dependent and abolished when high concentrations of echinacoside were applied (Fig. 3a and b, Fig. 4a), consistent with its influence on aggregate sizes (Fig. S5). One possible explanation was the desolvation of echinacoside with multiple hydroxyl groups when applied at higher concentrations, which might provide a substrate for the adhesion of bacteria similar to previously reported polymers^71^. Indeed, during experiments, we visibly observed substantial precipitation in 20 mM echinacoside solution in H_2_O (max solubility 45.39 mM) when it was incubated at room temperature. Nevertheless, future work will focus on understanding the mechanisms of this concentration-dependent manner that offer the basis for potential molecular modification and/or the development of new formulation and delivery strategies.

In conclusion, this study for the first time systematically investigated the expression profiles of c-di-GMP-related genes among various CF *P. aeruginosa* isolates in a pre-clinical model and discovered this antibiotic-potentiating function of echinacoside against *P. aeruginosa*. Our findings not only pave the way for developing novel combination therapy with clinical potential using echinacoside and tobramycin, but also reinforce the importance of using clinically relevant models for future drug discovery as stated by many researchers in recent years^3,22,72^.

## Materials and methods

### Strain and culture conditions

Bacterial strains and constructed plasmids in this study are listed in Table S2. The incubation time for each strain is listed in Table S4. Overnight cultures were grown in Lennox lysogeny broth (LB, Neogen, UK). Bacterial aggregates were grown in SCFM2 prepared as previously described^73^, while corresponding planktonic cultures for cyclic di-GMP ELISA assay and RNA extraction were grown in SCFM2 without salmon sperm DNA and mucin.

### Genetic manipulation in PAO1 and PA14

In-frame, seamless gene deletion on the PAO1 chromosome and complementary strains were constructed as previously described^74^. Primer pairs are listed in Table S6. To generate GFP-tagged PA14, pBK-miniTn7-gfp2 plasmid in *E. coli* DH5α was conjugated into PA14 through four-parental mating as previously described^74^. Colonies grown on PIA containing 100 μg/mL gentamicin were subjected to the fluorescent microscope for confirmation.

### Whole genome sequencing, RNA extraction and RT-qPCR

Genomic DNA from different isolates was obtained from pelleted overnight cultures using the Wizard Genomic DNA purification kit (Promega) following the manufacturer’s instructions. The quality of extracted DNA was measured using BioDrop μLITE (BioDrop, Cambridge, UK) before library preparation (350 bp) and Illumina sequencing (NovaSeq PE150, Novogene, UK). Contigs were assembled using the QIAGEN CLC genomic workbench. Accession numbers are listed in Table S2. Each c-di-GMP metabolism-related gene in each strain was identified *via* BLASTN (Table S3).

For total RNA extraction, overnight cultures of different strains were washed twice in 0.9% NaCl before inoculated into SCFM2 (for aggregate culture) and SCFM2 w/o DNA and mucin (for planktonic culture) at a final concentration of 1×10^7^ CFU/mL. Aggregates were grown statically in U-bottom 96 well plates (200 µL each well) in microaerophilic incubator (3% O_2_, 5% CO_2_) at 37. Planktonic cells were grown in 15 mL falcon tubes (3 mL each) with limited access to O_2_ (lid screwed tight) at 37 with vigorous shaking (250 rpm). After the desired incubation time for different strains (Table S4), 1200 µL combined bacterial aggregates or 600 µL planktonic cells were mixed 1:1 with RNAlater (Invitrogen, Thermo Fisher). Samples were then pelleted at 5000×*g* at 4 for 15 mins and re-suspended in 200 µL of freshly prepared 25 mg/mL lysozyme (Thermo Fisher) dissolved in 1×Tris-EDTA buffer (Fisher). The total RNA in each sample was extracted using QIAGEN RNeasy minikit and RNase-free DNase set following the manufacturer’s instructions. The residual gDNA in each sample was removed by TURBO DNA-free™ Kit (Invitrogen, Thermo Fisher) following the manufacturer’s instructions. The quality and quantity of RNA in each sample were measured by BioDrop μLITE.

Freshly extracted RNA (500 ng per 20µL reaction) was transcribed into cDNA using High-Capacity cDNA Reverse Transcription Kit (Applied Biosystems, Thermo Fisher). Such cDNA samples were applied as templates for RT-qPCR using 2× GoTaq® qPCR Master Mix (Promega). The reactions were performed on the CFX96^TM^ system (BIO-RAD) with the following PCR program: 95°C for 3 min; 40 cycles of 95°C for 15 s, Tm for 30 s and 72°C for 15s; plate read and melt curve generation. RT-qPCR primers for each gene were designed using Clone Manager 8, and the sequence of each primer was 100% conserved in all tested strains (Table S6). The specificity of each primer pair was tested by PCR, and the optimal annealing temperature was tested *via* thermal gradient RT-qPCR using PAO1 cDNA as the template. The stability of 7 frequently used housekeeping genes (*ampC*, *oprD*, *rpsL*, *fabD*, *rpoS*, *gyrA*, *proC*) in *P. aeruginosa* was tested using RefFinder^75^, which integrates the calculation from 4 major computational programs (geNorm, Normfinder, BestKeeper, and the comparative *Δ*-Ct method). An appropriate weight to each gene was assigned, and the geometric mean of their weights was calculated for the comprehensive ranking (supplemental RT-qPCR dataset; Fig. S1d). The relative transcript levels of target genes were then quantified by being normalized against *proC* showing the highest stability. Heatmap was generated by an online Web tool: https://biit.cs.ut.ee/clustvis/^76^.

### Virtual screening and molecular docking

The software used for virtual screening and 3-D mapping were Schrödinger Maestro 11.4 and PyMOL, respectively. For protein structure preparation, the crystal structure of SiaD was obtained from http://www.rcsb.org/ (PDB ID: 7E6G). 21495 bioactive compounds, including natural products, enzyme inhibitors, receptor ligands, and drugs from MCE Bioactive Compound Library (MedChemExpress, HY-L001V), were prepared using the LigPrep module. Compounds were initially docked by the Glide high-throughput virtual screening (HTVS) mode. The top 15% ranked compounds were chosen and redocked by the Glide standard precision (SP) scoring mode. Then the top 15% ranked compounds from this selected group were subjected to another round of docking by the extra precision (XP) scoring mode.

### Evaluation of the antimicrobial susceptibility to tobramycin and selected compounds

The Minimum inhibitory concentration (MIC) values of tobramycin (MedChemExpress), ciprofloxacin hydrochloride monohydrate (MedChemExpress), ceftazidime (MedChemExpress) and meropenem (Fresenius Kabi, Belgium) were determined in Mueller-Hinton broth (MHB, Neogen, UK) using EUCAST microdilution method. The tested concentration range of each antibiotic was 0.0625 µg/mL - 512 µg/mL. Pre-established aggregates of different *P. aeruginosa* strains were grown in SCFM2 in U-shape 96 well plates in a microaerophilic incubator after desired periods (Table S4). The efficacy of each antibiotic against pre-established aggregates was determined within the working concentration range of 4×MIC - 512 µg/mL after 18-hr treatment. Treated and untreated samples were collected into 2 mL reinforced tubes and subjected to bead ruptor 24 elite (OMNI, USA) for aggregate disruption (4000 rpm, 2 mins), and CFU numbers were determined by plating. Concentrations of different antibiotics where at least 1-log reduction of CFU can be observed compared to untreated samples but did not completely eradicate aggregates were used for further tests.

Different concentrations of echinacoside were applied either alone or with different antibiotics at the abovementioned concentrations. Echinacoside was dissolved in H_2_O at a concentration of 20 mM as stock solution. The bactericidal effect of compounds alone and the effect of combination treatments were determined by plating and CFU count as abovementioned.

### Quantification of intracellular c-di-GMP levels

Intracellular c-di-GMP quantification for aggregates and planktonic cultures was accomplished by c-di-GMP ELISA kit (Cayman, USA). To compare the difference between aggregated and planktonic samples, bacteria were cultured under identical conditions as those cultured for RNA extraction. To assess the effect of echinacoside on c-di-GMP, different concentrations of the compound were added together with bacteria diluted in SCFM2 (1×10^7^ CFU/mL). After desired incubation, 1200 µL aggregate samples and 600 µL planktonic samples were collected and immediately transferred to pre-chilled centrifuge tubes kept on ice. 100 µL of each sample was spared for CFU determination. Samples were centrifuged at 5000×*g* for 15 mins at 4, and supernatant was discarded. Pellets were then completely dissolved in 200 µL B-PER™ bacterial protein extraction reagent (Thermo Fisher) for 15 mins at room temperature prior to centrifugation (15000×*g*, 10 mins). Supernatants from aggregate and planktonic samples were 3-fold and 6-fold diluted into B-PER solution, respectively, for ELISA assay, and 2-fold diluted into MQ H_2_O to measure total protein concentration. Diluted samples were subjected to c-di-GMP ELISA kit and Pierce™ 660nm protein assay reagent (Thermo Fisher) following manufacturer’s instructions. 2 mg/mL bovine serum albumin (BSA, Thermo Fisher) was used to determine standard curves for protein quantification. Total c-di-GMP was then normalized to CFU or total protein.

### 3-D lung epithelial model infection assay and cytotoxicity assay

The 3-D *in vivo*-like lung model was established from the human alveolar epithelial cell line A549 using GTSF-2 medium (HyClone, Logan, UT, US) as previously described^77^. On the day of infection, an equal number of 3-D A549 cells were transferred into each well in flat-bottom 24 well plates to achieve ∼2.5 × 10^5^ cells/well. Mid-log phase GFP-tagged PA14 (for microscopic analysis) or PA14 (for cytotoxicity assay) bacterial suspensions grown in LB were washed twice in GTSF-2 medium without FBS and inoculated into each well containing 3-D A549 cells at a multiplicity of infection (MOI) of 100:1. After 4-h static co-culture in a microaerophilic incubator at 37 to allow for bacterial adherence, cells were washed twice with GTSF-2 without FBS to discard planktonic or loosely attached PA14. PA14-infected 3-D A549 cells were treated with tobramycin, echinacoside, or both together for 18 hrs in a microaerophilic incubator statically. The viability of A549 cells with and without different treatments were quantified using ‘intracellular’ lactate dehydrogenase (LDH) assay as previously described^78^.

### Microscopic analysis

For observing bacterial aggregates, overnight cultures of PAO1 WT and mutants were washed twice in 0.9% NaCl and diluted into SCFM2. To compare the size of aggregates formed by different isogenic mutants, diluted bacteria suspensions at a final concentration of 1×10^5^ CFU/mL in SCFM2 were inoculated into flat-bottom 48 well plates (1 mL per well) and incubated in a microaerophilic incubator statically for 24 hrs. To assess the effect of selected compounds on auto-aggregation, different concentrations of compounds were added together with PAO1 WT diluted samples (1×10^5^ CFU/mL) and incubated statically for 6 hrs. To observe aggregates (grown statically) and planktonic cells (grown with 250 rpm shaking), overnight cultures were diluted to a final concentration of 1×10^7^ CFU/mL in SCFM2 or SCFM2 without mucin and DNA. After incubation, samples were harvested carefully with wide-orifice tips (Finntip, Thermo Fisher) from the middle layer of bacteria suspension in each well (aggregate)/each tube (planktonic) and 4-fold (aggregate)/12-fold (planktonic) diluted into SCFM2 without DNA and mucin. 200 µL such diluted samples were inoculated into each well in uncoated Ibidi 96 well plate (Ibidi, USA) and subjected to EVOS FL Auto microscope (Thermo Fisher). Micrographs were obtained using 20× objective with transmitted light. The gap between each z-stack was 0.488 µm. The size of each aggregate in image stacks was quantified by Fiji as previously described^74^.

For observing the overall morphology/integrity of 3-D lung epithelial cells and attached GFP-tagged PA14, micrographs were obtained using 10× objective with both transmitted light and GFP fluorescent light cube (470 nm excitation, 525 nm emission).

## Acknowledgment

The authors would like to thank Dr. Andrea Sass for the discussion of RT-qPCR set-up and data analysis, Lisa Ostyn for the help with 3-D cell culture and Amber de Craemer for the help with bactericidal tests. *P. aeruginosa* clinical isolates (BS and OS) collected from Ghent University Hospital were generously provided by Dr. Sara Van den Bossche. This project was supported by MSCA-IF-EF-ST (Grant #101023767) to YMC.

## Author contributions

YMC and TC conceptualized the study. YMC designed experimental methods with input from TC and AC. YMC carried out the laboratory experiments and analysed the data. YMC and TC wrote the manuscript, with edition from AC. All authors contributed to the article and approved the submitted version.

