## supplemental for "Identification of echinacoside as a tobramycin potentiator against *Pseudomonas aeruginosa* aggregates"

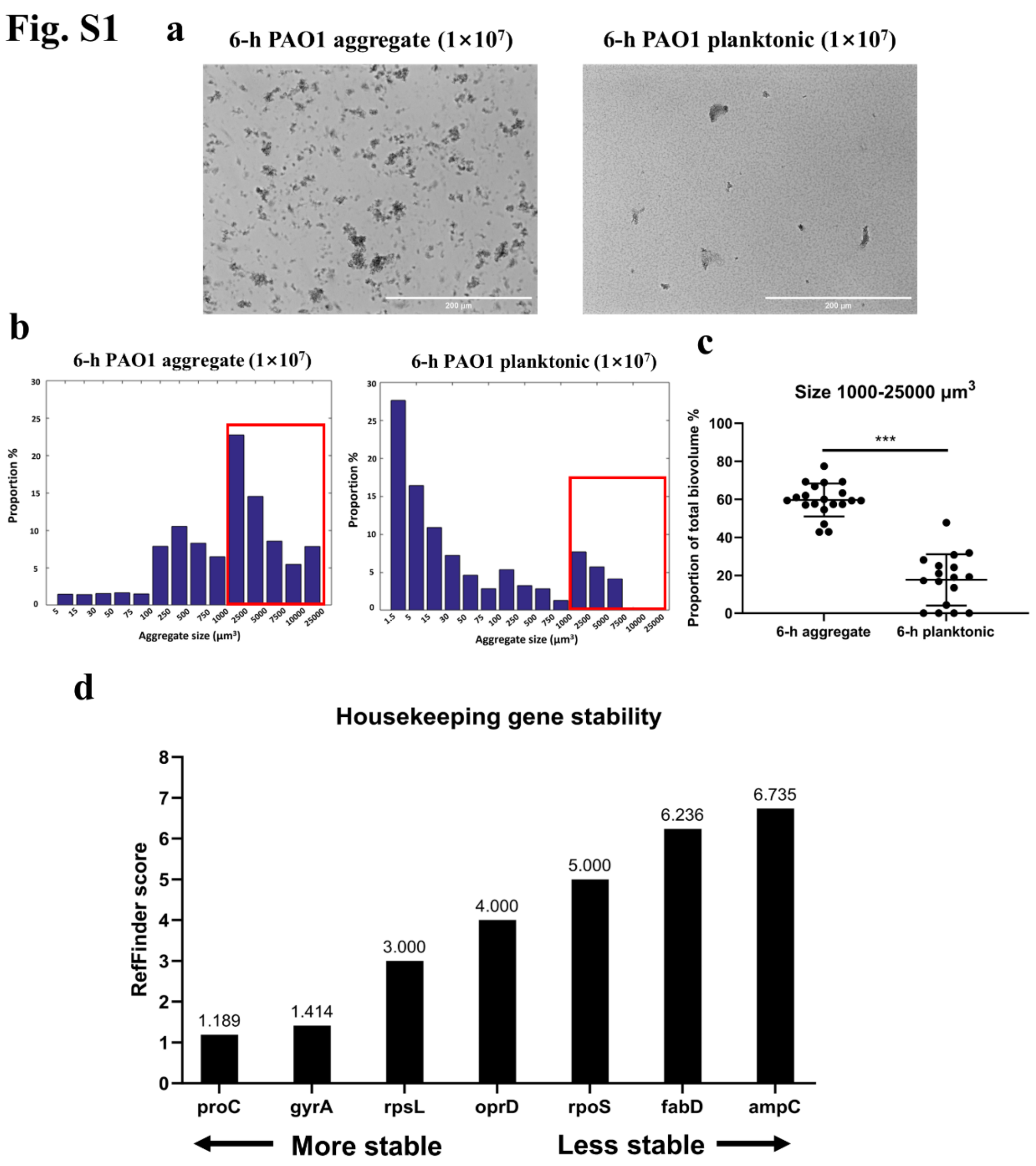

**Fig. S1 (a)** Representative micrographs of *P. aeruginosa* PAO1 WT aggregates (in SCFM2) and planktonic cells (in SCFM2 without mucin and DNA) grown for 6 hrs, with an initial inoculum size of $1\times10^7$ CFU/mL. Scale bar = 200 μm. **(b)** Distribution of the size of 6-h PAO1 WT aggregates and planktonic cells. Histograms represent the mean proportion value of all samples collected from 3 individual experiments (n=3), where micrographs were obtained from at least 3 random locations in each sample. Numerical data and standard deviations are shown in Table S1. **(c)** Proportion of aggregates with sizes ranging from 1000 to 25000 $\mu m^3$ in total biovolume. ***, $p<0.001$ (Student's t-test). **(d)** Stability of expression of 7 candidate reference genes for qPCR normalization as determined with RefFinder. The stability index of each gene is shown above each bar.

**Fig. S2**

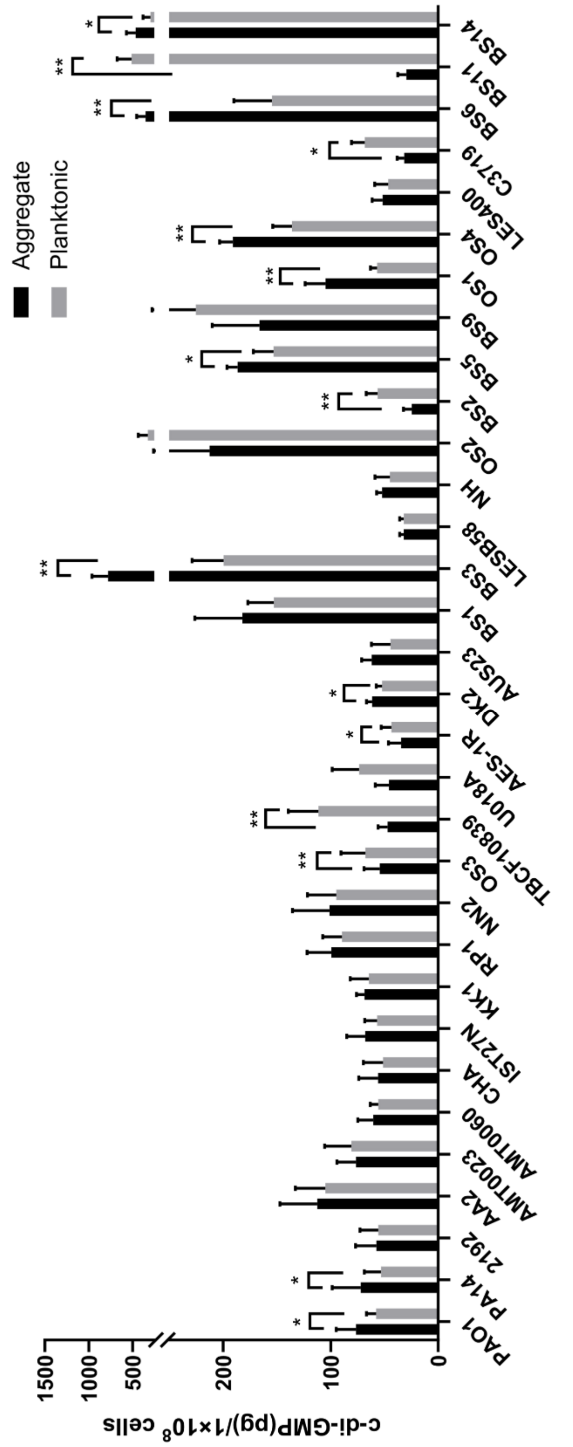

**Fig. S2** Intracellular c-di-GMP levels for 32 *P. aeruginosa* strains grown as aggregates (in SCFM2) or planktonic cells (in SCFM2 without DNA and mucin). Intracellular c-di-GMP concentrations (as determined by ELISA) were normalized to CFU counts (determined by plating). Data were acquired from 3 independent experiments with 2 technical replicates and the mean values were shown. Error bars indicate standard deviation. *, $p<0.05$; **, $p<0.01$ (Ratio paired t-test between two groups).

**Fig. S3**

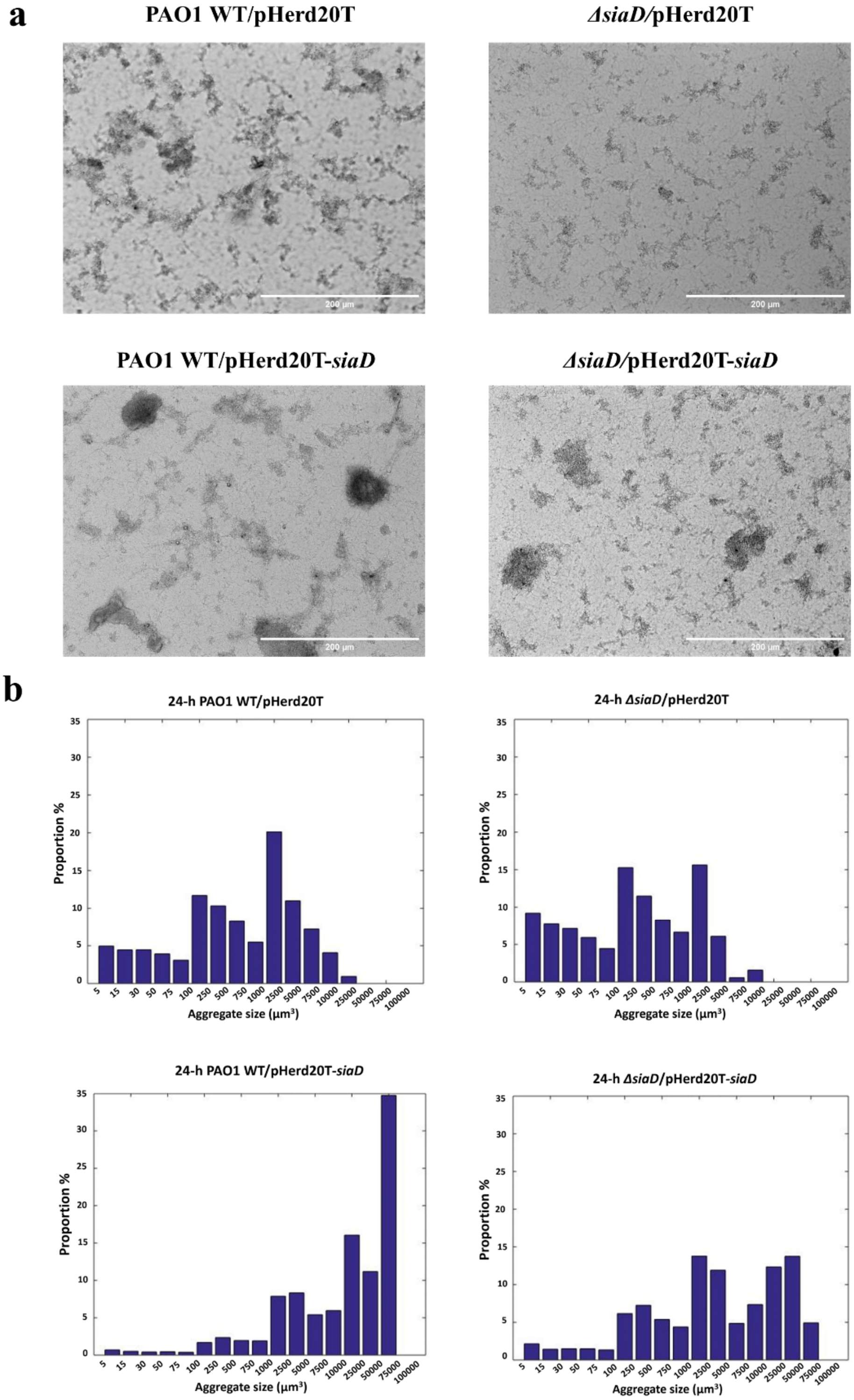

**Fig. S3 (a)** Representative micrographs of PAO1 WT/pHerd20T, *ΔsiaD*/pHerd20T-*siaD,* PAO1 WT/pHerd20T-*siaD* and *ΔsiaD*/pHerd20T-*siaD* aggregates grown for 24 h in SCFM2. Scale bar = 200 μm. Plasmids were maintained by adding 200 μg/mL carbenicillin and expression was induced by 0.5% (v/v) arabinose. **(b)** Distribution of the size of 24-h PAO1 WT/pHerd20T, *ΔsiaD*/pHerd20T-*siaD,* PAO1 WT/pHerd20T-*siaD* and *ΔsiaD*/pHerd20T-*siaD* aggregates grown in SCFM2. The biovolume ($\mu m^3$) of single cells and aggregates were grouped into 17 categories. Histograms represent the mean proportion value of all samples collected from 3 individual experiments (n=3), where micrographs were obtained from at least 3 random locations in each sample. Numerical data of mean proportions and standard deviation are shown in Table S1.

**Fig. S4**

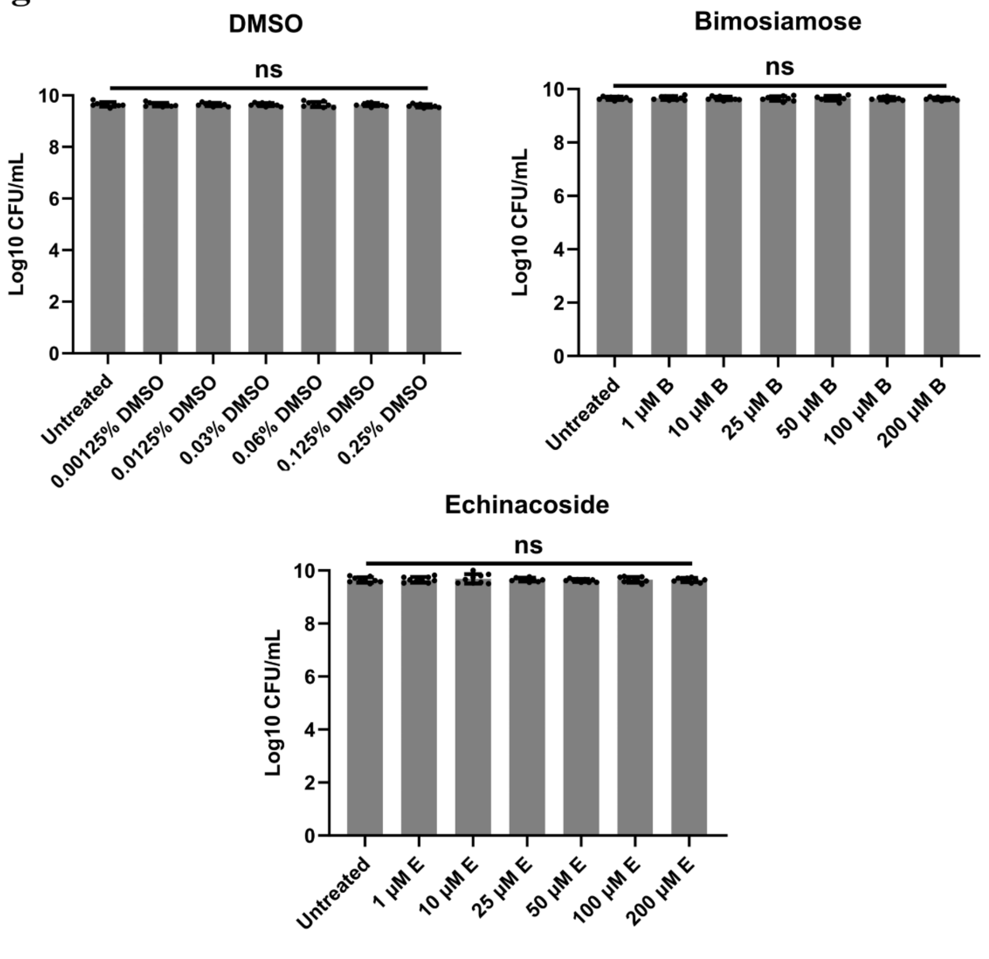

**Fig. S4** Bactericidal tests for DMSO (solvent for bimosiamose), bimosiamose, and echinacoside against 6-h pre-established PAO1 aggregates grown in SCFM2. Data are expressed as the mean number of CFU remaining after an additional 18-h incubation (3 independent experiments with 3 technical replicates. No statistically significant difference was found comparing untreated and treated samples (One-way ANOVA).

Fig. S5

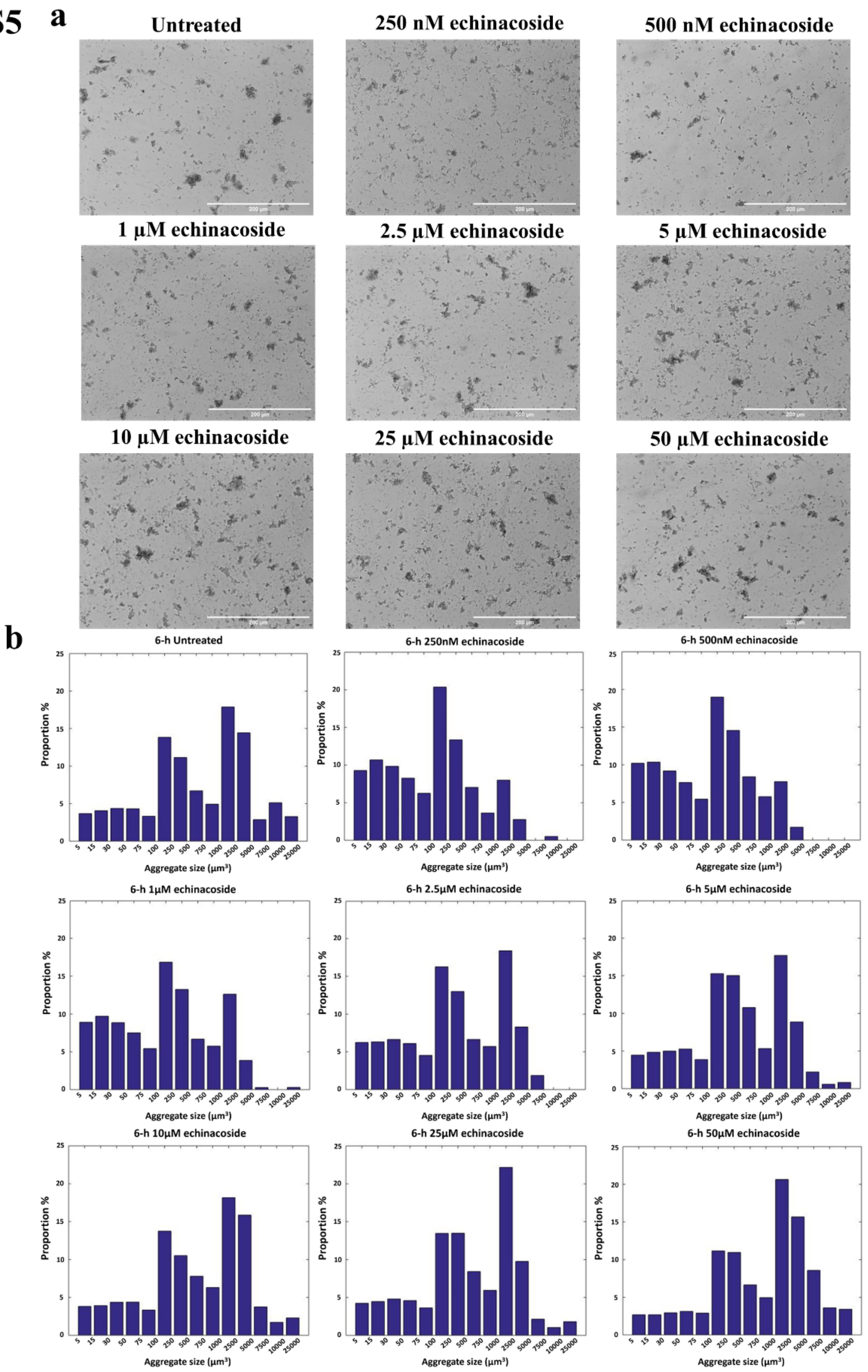

**Fig. S5 (a)** Representative micrographs of 6-h *P. aeruginosa* PAO1 aggregates grown in SCFM2 treated with different concentrations of echinacoside. Scale bar = 200 µm. **(b)** Distribution of the size of 6-h PAO1 aggregates grown in SCFM2 treated with different concentrations of echinacoside. The biovolume ($\mu m^3$) of single cells and aggregates were grouped into 14 categories. Histograms represent the mean proportion value of all samples collected from 3 individual experiments (n=3), where micrographs were obtained from at least 3 random locations in each sample. Numerical data of mean proportions and standard deviation are shown in Table S1.

**Fig. S6**

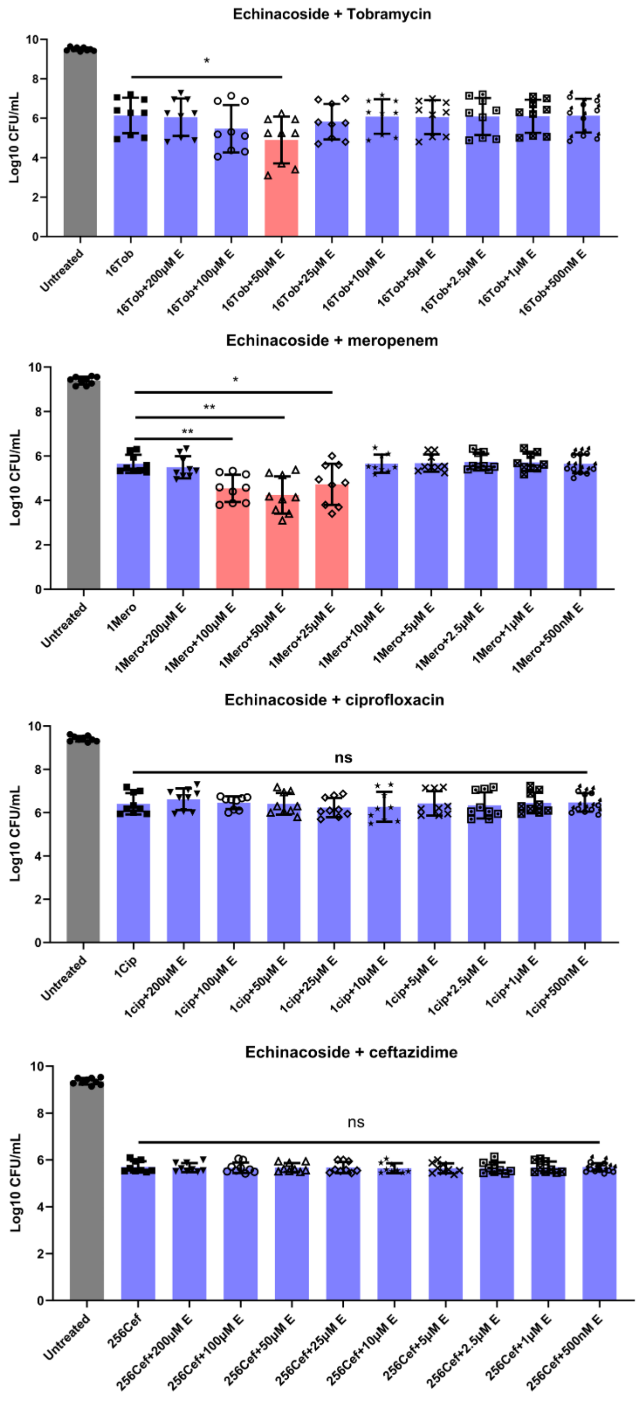

**Fig. S6** Antimicrobial activity of the combination of echinacoside with tobramycin, meropenem, ciprofloxacin, and ceftazidime, respectively, against 6-h pre-established *P. aeruginosa* PAO1 aggregates grown in SCFM2. Data are expressed as the mean number of CFU remaining after an additional 18-h incubation (3 independent experiments with 3 technical replicates; error bars indicate standard deviation). Red bars highlighted the successful combination treatment that potentiated the efficacy of corresponding antibiotics. *, p<0.05; **, p<0.01 (Student's t-test).

**Fig. S7**

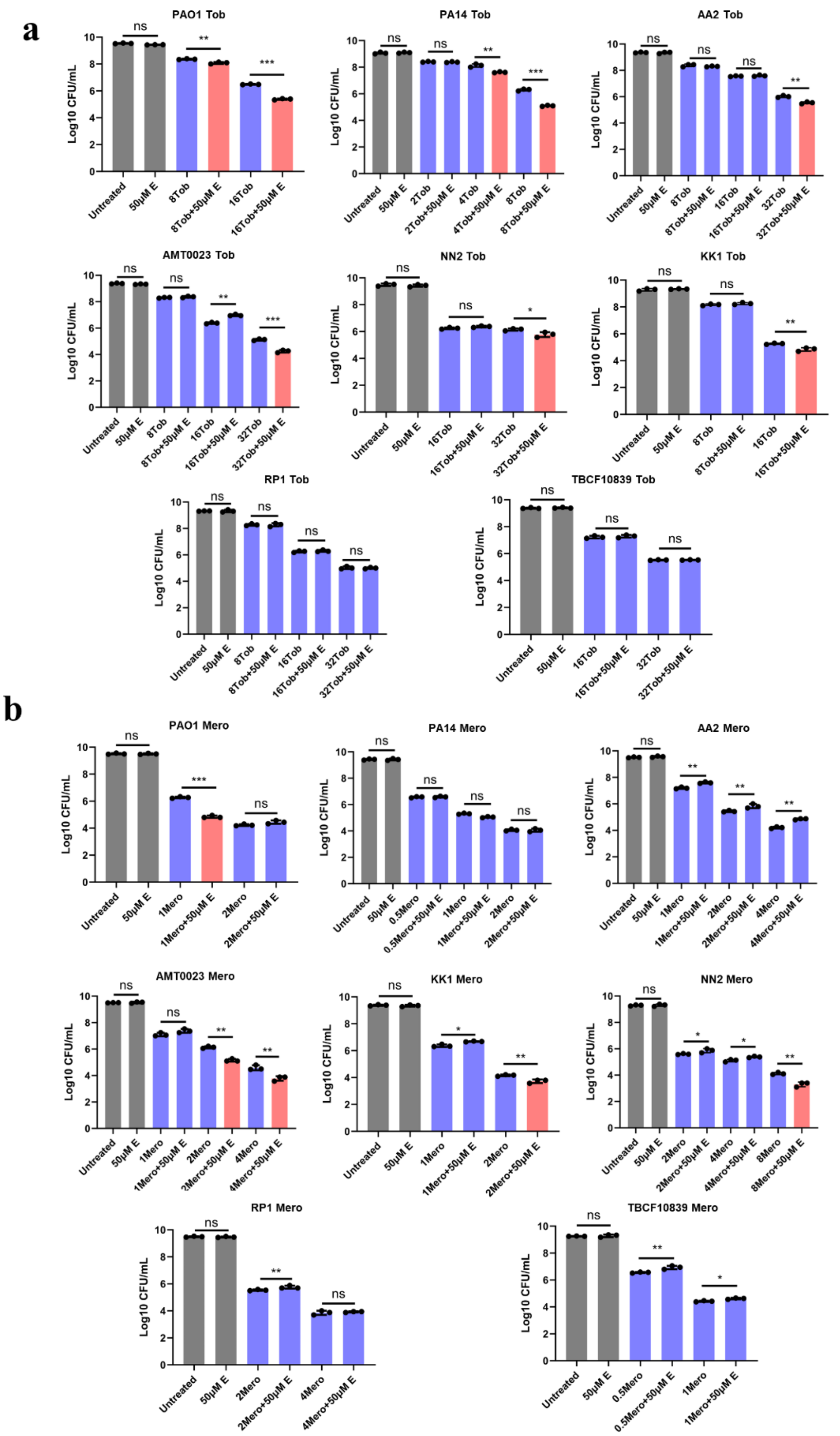

**Fig. S7** Efficacy test for the combination treatment of 50 μM echinacoside with different concentrations of **(a)** tobramycin or **(b)** meropenem against pre-established aggregates formed by 8 different *P. aeruginosa* strains in SCFM2. Data are expressed as the mean number of CFU remaining after an additional 18-h incubation (3 independent experiments with 3 technical replicates; error bars indicate standard deviation). Red bars highlighted the successful combination treatment that potentiated the efficacy of corresponding antibiotics. *, p<0.05; **, p<0.01; ***, p<0.001 (Student's t-test).

**Fig. S8**

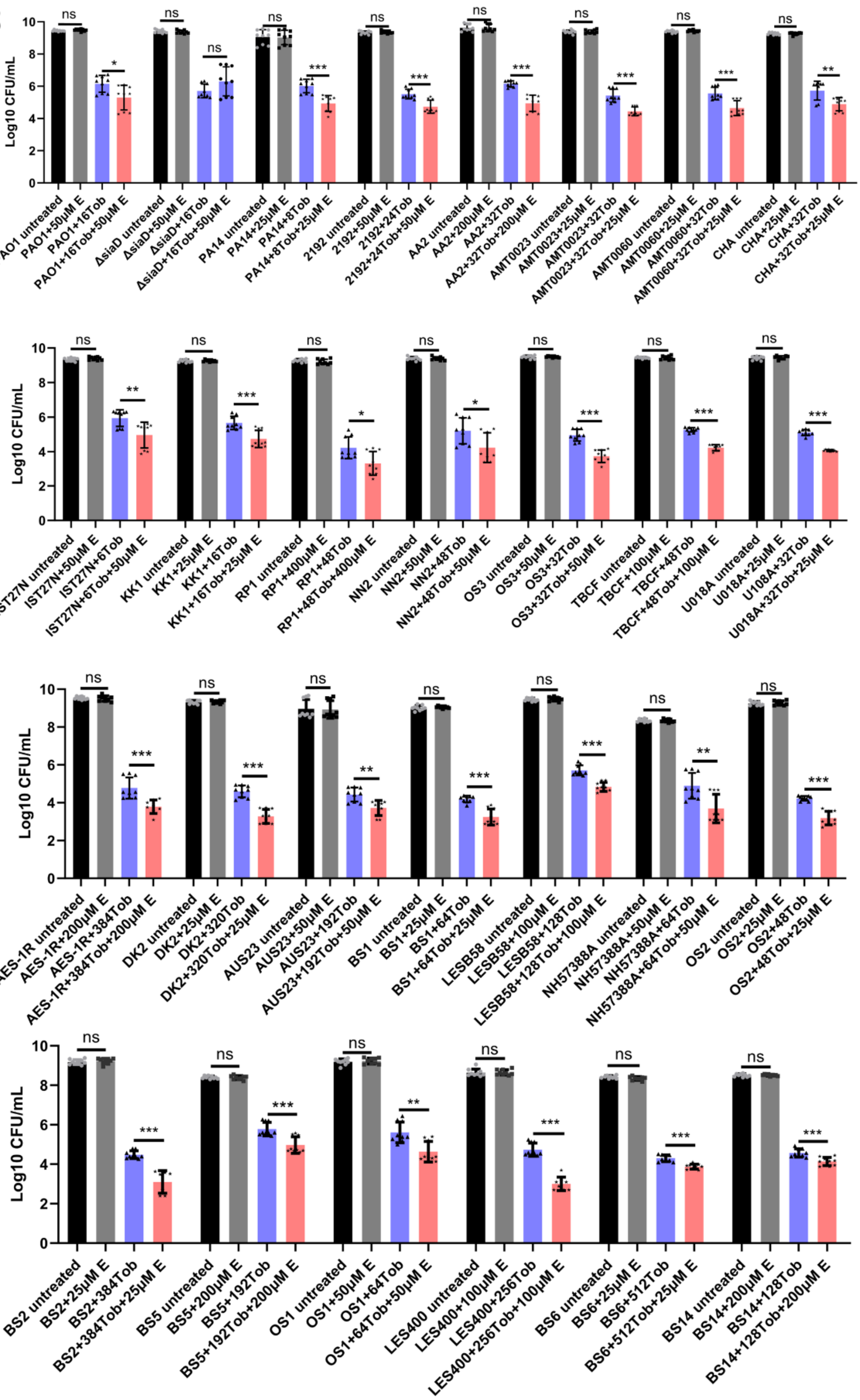

**Fig. S8** The optimal dosages for successful combination treatments where echinacoside potentiated the efficacy of tobramycin against pre-established aggregates grown in SCFM2 for 28 *P. aeruginosa* strains. Data are expressed as the mean number of CFU remaining after an additional 18-h incubation (3 independent experiments with 3 technical replicates; error bars indicate standard deviation). *, p<0.05; **, p<0.01; ***, p<0.001 (Student's t-test).

### Fig. S9

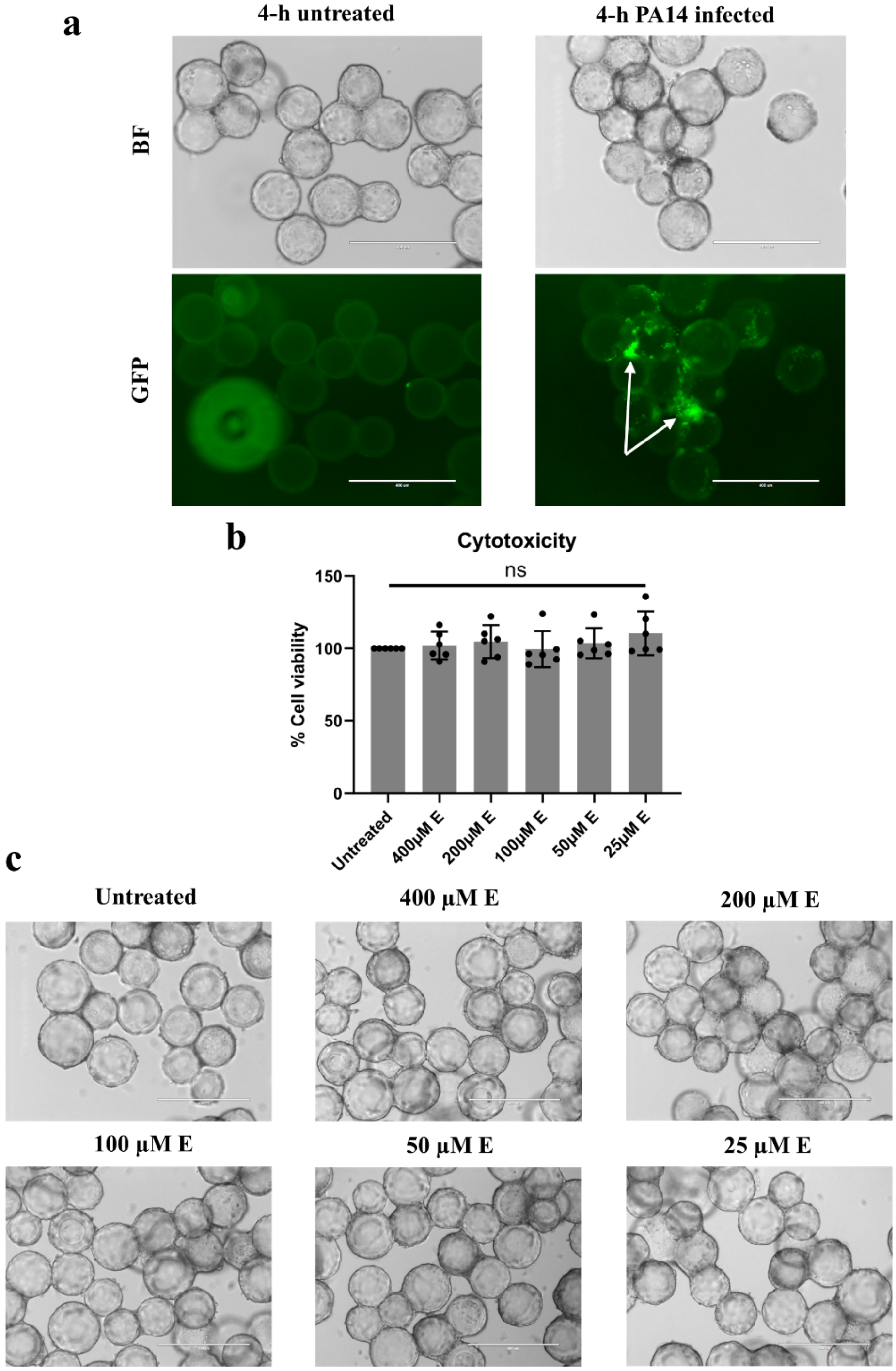

**Fig. S9** **(a)** Representative transmission light micrographs (upper panel) of 3-D A549 cells attached to/detached from collagen-treated carrier beads and fluorescent micrographs (lower panel) of GFP-tagged PA14 attached to A549 cells. After 4-h infection, PA14 aggregates (highlighted with white arrows) were found attached to A549 cells with strong fluorescent signals. Healthy cells in the untreated group showed a smooth morphology and remained firmly attached to beads, while cells infected with PA14 were stressed (rough and patchy morphology at the surface of beads). Scale bar = 400 μm. **(b)** Cytotoxicity test for different concentrations of echinacoside. The viability of 3-D A549 cells challenged with different concentrations of echinacoside for 18 hrs was measured by LDH assay and normalized to untreated groups. Data were acquired from 6 independent experiments with 4 technical replicates. No statistically significant difference was found comparing untreated and treated cells (One-way ANOVA). (c) Representative transmission light micrographs of 3-D A549 cells attached to collagen-treated carrier beads challenged with different concentrations of echinacoside. Healthy cells in all groups showed a smooth morphology and remained firmly attached to beads without difference in attachment patterns. Scale bar = 400 μm.

Table S1. Mean fraction (%) of aggregates within each size category in total biovolume. SD, standard deviation. N=3

| 6-h aggregates (initial inoculum $1\times10^5$ CFU/mL) | | | | | | | | | | | | | | | | | | |
|---|---|---|---|---|---|---|---|---|---|---|---|---|---|---|---|---|---|---|
| Size µm$^3$ | Untreated | | 250 nM E | | 500 nM E | | 1 µM E | | 2.5 µM E | | 5 µM E | | 10 µM E | | 25 µM E | | 50 µM E | |
| | mean | SD | mean | SD | mean | SD | mean | SD | mean | SD | mean | SD | mean | SD | mean | SD | mean | SD |
| 5-15 | 3.67 | 1.2 | 9.29 | 2.93 | 10.21 | 4.5 | 9.21 | 4.31 | 6.24 | 2.28 | 4.46 | 1.68 | 3.81 | 1.44 | 4.24 | 1.29 | 2.69 | 0.95 |
| 15-30 | 4.07 | 1.78 | 10.69 | 2.6 | 10.34 | 2.68 | 9.87 | 3.9 | 6.31 | 1.83 | 4.83 | 1.43 | 3.92 | 1.61 | 4.46 | 1.67 | 2.69 | 1.12 |
| 30-50 | 4.37 | 1.79 | 9.85 | 1.82 | 9.18 | 2.61 | 8.81 | 3.16 | 6.64 | 2.15 | 5 | 0.95 | 4.37 | 1.66 | 4.81 | 1.4 | 2.94 | 1.17 |
| 50-75 | 4.32 | 1.34 | 8.26 | 1.27 | 7.65 | 1.88 | 7.59 | 2.38 | 6.11 | 1.88 | 5.26 | 0.78 | 4.39 | 1.44 | 4.6 | 1.17 | 3.13 | 1.51 |
| 75-100 | 3.32 | 0.76 | 6.25 | 0.84 | 5.44 | 1.83 | 5.5 | 1.74 | 4.53 | 2.22 | 3.86 | 0.78 | 3.34 | 0.62 | 3.64 | 0.94 | 2.89 | 1.38 |
| 100-250 | 13.84 | 3.29 | 20.37 | 3.11 | 18.99 | 3.58 | 16.69 | 3.95 | 16.26 | 3.6 | 15.28 | 1.3 | 13.75 | 2.65 | 13.46 | 2.2 | 11.16 | 7.42 |
| 250-500 | 11.15 | 3.51 | 13.36 | 2.82 | 14.56 | 4.06 | 12.83 | 2.6 | 12.99 | 3.37 | 15.03 | 2.73 | 10.52 | 1.72 | 13.48 | 2.06 | 10.96 | 8.72 |
| 500-750 | 6.72 | 2.45 | 7.03 | 1.76 | 8.41 | 4.41 | 6.82 | 2.97 | 6.65 | 1.73 | 10.77 | 1.84 | 7.79 | 2.33 | 8.43 | 2.16 | 6.65 | 0.98 |
| 750-1000 | 4.93 | 1.85 | 3.63 | 2.11 | 5.77 | 4.12 | 5.62 | 2.73 | 5.71 | 3.28 | 5.33 | 1.45 | 6.31 | 1.75 | 5.96 | 1.42 | 4.96 | 1.59 |
| 1000-2500 | 17.89 | 5.23 | 8 | 6.1 | 7.76 | 5.11 | 12.12 | 9 | 18.38 | 9.19 | 17.7 | 4.92 | 18.17 | 3.42 | 22.18 | 3.26 | 20.65 | 17.71 |
| 2500-5000 | 14.44 | 9.06 | 2.77 | 3.84 | 1.71 | 3.03 | 4.2 | 5.29 | 8.29 | 7.39 | 8.86 | 4.75 | 15.87 | 6.17 | 9.77 | 3.08 | 15.68 | 13.59 |
| 5000-7500 | 2.87 | 3.86 | 0 | 0 | 0 | 0 | 0.44 | 1.09 | 1.88 | 3.85 | 2.21 | 2.69 | 3.76 | 4.21 | 2.14 | 2.96 | 8.57 | 18.81 |
| 7500-10000 | 5.13 | 5.09 | 0.51 | 1.97 | 0 | 0 | 0 | 0 | 0 | 0 | 0.59 | 1.76 | 1.71 | 2.67 | 1.04 | 2.16 | 3.61 | 10.35 |
| 10000-25000 | 3.28 | 4.37 | 0 | 0 | 0 | 0 | 0.3 | 1.33 | 0 | 0 | 0.82 | 2.46 | 2.29 | 5.31 | 1.81 | 3.65 | 3.41 | 17.51 |

| 24-h aggregates (initial inoculum $1\times10^5$ CFU/mL) | | | | | | | | | | | | | | | | | | |
|---|---|---|---|---|---|---|---|---|---|---|---|---|---|---|---|---|---|---|
| Size µm$^3$ | PAO1 WT (1) | | *ΔwspR* | | PAO1 WT (2) | | *Δpa5442* | | *ΔsiaD* | | WT/pHerd20T | | WT/pHerd20T-*siaD* | | *ΔsiaD*/pHerd20T | | *ΔsiaD*/pHerd20T-*siaD* | |
| | mean | SD | mean | SD | mean | SD | mean | SD | mean | SD | mean | SD | mean | SD | mean | SD | mean | SD |
| 5-15 | 1.49 | 1.04 | 1.65 | 0.99 | 1.95 | 1.22 | 1.76 | 1.09 | 16.78 | 9.6 | 4.95 | 4.01 | 0.71 | 0.22 | 9.2 | 5.05 | 2.14 | 1.17 |
| 15-30 | 1.37 | 1.23 | 1.51 | 0.86 | 1.86 | 1.08 | 1.63 | 0.82 | 12.19 | 4.32 | 4.45 | 3.07 | 0.52 | 0.19 | 7.74 | 2.7 | 1.43 | 0.57 |
| 30-50 | 1.3 | 1.09 | 1.49 | 0.98 | 1.78 | 0.99 | 1.6 | 0.67 | 11.28 | 4.04 | 4.46 | 2.85 | 0.42 | 0.15 | 7.13 | 2.19 | 1.5 | 0.62 |
| 50-75 | 1.36 | 0.95 | 1.36 | 0.86 | 1.73 | 1.03 | 1.75 | 0.84 | 9.64 | 3.57 | 3.93 | 2.03 | 0.46 | 0.24 | 5.92 | 1.66 | 1.49 | 0.54 |
| 75-100 | 1.17 | 0.8 | 1.33 | 0.68 | 1.47 | 0.85 | 1.44 | 0.74 | 6.12 | 2.67 | 3.08 | 1.29 | 0.39 | 0.19 | 4.45 | 1.17 | 1.34 | 0.5 |
| 100-250 | 5.66 | 3.62 | 5.53 | 2.18 | 6.17 | 2.99 | 6.95 | 3.39 | 23.29 | 7.9 | 11.71 | 3.51 | 1.71 | 0.76 | 15.28 | 2.66 | 6.15 | 2.29 |
| 250-500 | 6.77 | 4.12 | 6.37 | 1.95 | 6.21 | 2.57 | 7.97 | 3.71 | 11.51 | 8.35 | 10.32 | 2.7 | 2.36 | 1 | 11.49 | 4.28 | 7.25 | 2.38 |
| 500-750 | 5.06 | 3.04 | 4.93 | 1.69 | 4.45 | 1.05 | 6.4 | 2.29 | 4 | 4.02 | 8.26 | 2.26 | 1.96 | 0.75 | 8.28 | 3.09 | 5.37 | 1.62 |
| 750-1000 | 4.09 | 2.58 | 4.8 | 2.25 | 3.32 | 1 | 5.28 | 2.67 | 1.87 | 3.13 | 5.49 | 1.94 | 1.91 | 0.9 | 6.64 | 3.32 | 4.39 | 1.23 |
| 1000-2500 | 15.56 | 6.07 | 17.92 | 5.14 | 14.6 | 5.28 | 19.25 | 5.77 | 2.73 | 4.93 | 20.12 | 5.83 | 7.88 | 3.68 | 15.64 | 6.73 | 13.78 | 6.61 |
| 2500-5000 | 14.45 | 6.18 | 16.92 | 5.06 | 12.42 | 3.97 | 17.68 | 6.6 | 0.58 | 2.24 | 11 | 6.77 | 8.34 | 2.29 | 6.08 | 6.71 | 11.91 | 5.79 |
| 5000-7500 | 7.15 | 4.15 | 12.06 | 7.23 | 9.44 | 5.78 | 11.43 | 7.1 | 0 | 0 | 7.21 | 8.25 | 5.41 | 3.75 | 0.57 | 2.21 | 4.86 | 4.36 |
| 7500-10000 | 4.43 | 4.66 | 5.69 | 5.56 | 6.49 | 4.1 | 3.07 | 3.34 | 0 | 0 | 4.09 | 4.73 | 5.96 | 5.29 | 1.58 | 4.24 | 7.35 | 2.87 |
| 10000-25000 | 18.71 | 11.85 | 14.17 | 9.76 | 16.88 | 7.88 | 11.7 | 9.86 | 0 | 0 | 0.94 | 2.5 | 16.06 | 6.54 | 0 | 0 | 12.34 | 9.08 |
| 25000-50000 | 8.43 | 11.84 | 4.26 | 8.47 | 11.22 | 8.32 | 2.1 | 3.25 | 0 | 0 | 0 | 0 | 11.18 | 12.43 | 0 | 0 | 13.75 | 13.08 |
| 50000-75000 | 3.01 | 8.32 | 0 | 0 | 0 | 0 | 0 | 0 | 0 | 0 | 0 | 0 | 34.75 | 18.01 | 0 | 0 | 4.92 | 13.81 |
| 75000-100000 | 0 | 0 | 0 | 0 | 0 | 0 | 0 | 0 | 0 | 0 | 0 | 0 | 0 | 0 | 0 | 0 | 0 | 0 |

| 6-h aggregates vs planktonic (initial inoculum $1\times10^7$ CFU/mL) | | | | |
|---|---|---|---|---|
| Size µm$^3$ | aggregate | | planktonic | |
| | mean | SD | mean | SD |
| **1.5-5** | N/A | N/A | 27.64 | 10.31 |
| **5-15** | 1.44 | 0.48 | 16.44 | 4 |
| **15-30** | 1.39 | 0.43 | 10.92 | 3.94 |
| **30-50** | 1.52 | 0.48 | 7.26 | 2.73 |
| **50-75** | 1.64 | 0.49 | 4.65 | 2.25 |
| **75-100** | 1.48 | 0.5 | 2.82 | 1.56 |
| **100-250** | 7.89 | 3.04 | 5.38 | 3.21 |
| **250-500** | 10.56 | 2.59 | 3.23 | 1.55 |
| **500-750** | 8.31 | 2.36 | 2.81 | 2.14 |
| **750-1000** | 6.5 | 2.1 | 1.27 | 1.66 |
| **1000-2500** | 22.77 | 4.95 | 7.72 | 7.73 |
| **2500-5000** | 14.56 | 3.99 | 5.74 | 7.18 |
| **5000-7500** | 8.58 | 5.3 | 4.14 | 6.99 |
| **7500-10000** | 5.48 | 4.29 | 0 | 0 |
| **10000-25000** | 7.87 | 5.59 | 0 | 0 |
| **25000-50000** | 0 | 0 | 0 | 0 |

Table S2. Bacterial strains and plasmids used

| Clinical strains | | |
|---|---|---|
| *P. aeruginosa* isolates (accession number of genome sequence) | Description | Source or reference |
| PAO1 (NC_002516) | Wound isolate (Australia) | [1] |
| PA14 (NC_008463) | Burn wound isolate (US) | [2] |
| 2192 (NZ_CH482384) | CF isolate (US) | [3] |
| AA2 (MCMJ00000000) | CF isolate (Germany) | [4,5] |
| AES-1R (MCML00000000) | CF isolate (Australia) | [5,6] |
| AMT0023-30 (MCMI00000000) | CF isolate (US) | [5,7] |
| AMT0060-3 (MCMZ00000000) | CF isolate (US) | [5,7] |
| AUS23 (MCMN00000000) | CF isolate (Australia) | [5,8] |
| BS1 (PRJNA1072279) | CF isolate (Belgium) | This study |
| BS2 (PRJNA1072279) | CF isolate (Belgium) | This study |

| | | |
|---|---|---|
| BS3 (PRJNA1072279) | CF isolate (Belgium) | This study |
| BS5 (PRJNA1072279) | CF isolate (Belgium) | This study |
| BS6 (PRJNA1072279) | CF isolate (Belgium) | This study |
| BS9 (PRJNA1072279) | CF isolate (Belgium) | This study |
| BS11 (PRJNA1072279) | CF isolate (Belgium) | This study |
| BS14 (PRJNA1072279) | CF isolate (Belgium) | This study |
| C3719 (MCMM00000000) | CF isolate (UK) | [3,5] |
| CHA (MCMG00000000) | CF isolate (France) | [5,9] |
| DK2 (NC_018080) | CF isolate (Denmark) | [10] |
| IST27N (MCMW00000000) | CF isolate (Portugal) | [5,11] |
| KK1 (MCMB00000000) | CF isolate (Germany) | [4,5] |
| LESB58 (NC_011770) | CF isolate (UK) | [12] |
| LES400 (NZ_CP006982) | CF isolate (UK) | [13] |
| NH57388A (MCMT00000000) | CF isolate (Denmark) | [5,14] |
| NN2 (NZ_LT883143.1) | CF isolate (Germany) | [15,16] |
| OS1 (PRJNA1072279) | CF isolate (Belgium) | This study |
| OS2 (PRJNA1072279) | CF isolate (Belgium) | This study |
| OS3 (PRJNA1072279) | CF isolate (Belgium) | This study |
| OS4 (PRJNA1072279) | CF isolate (Belgium) | This study |
| RP1 (LNBU00000000) | CF isolate (Germany) | [17] |
| TBCF10389 (MCLZ00000000) | CF isolate (Germany) | [5,18] |
| U018A (MCMP00000000) | CF isolate (Australia) | [5,19] |
| *P. aeruginosa* PAO1 isogenic mutants | | |
| *ΔwspR* | *ΔwspR* deletion mutant | This study |
| *ΔsiaD* | *ΔsiaD* deletion mutant | This study |
| *Δpa5442* | *Δpa5442* deletion mutant | This study |
| PAO1 WT/pHerd20T | PAO1 carrying empty pHerd20T, $Cb^R$ | This study |
| *ΔsiaD*/pHerd20T | *ΔsiaD* mutant carrying empty pHerd20T, $Cb^R$ | This study |
| PAO1 WT/pHerd20T-*siaD* | PAO1 carrying pHerd20T with *siaD* ORF inserted, $Cb^R$ | This study |
| *ΔsiaD* /pHerd20T-*siaD* | *ΔsiaD* mutant carrying pHerd20T with *siaD* ORF inserted, $Cb^R$ | This study |
| GFP-labelled PA14 | | |
| PA14/pBK-miniTn7-gfp2 | Chromosomal labelled PA14 with pBK-miniTn7-gfp2. $Gm^R$, | This study |
| *E. coli* strains | | |
| HB101 | K-12/B hybrid; $Sm^r$ *recA thi pro leu hsdRM*$^+$ | [20] |
| DH5α | $F^-$ ϕ80dlacZΔM15 Δ(lacZYA-argF)*U169 deoR recA1 endA1 hsdR17*($r_K^-$ $m_K^+$) *phoA supE44 λ*$^-$ *thi-1 gyrA96 relA1* | [21] |
| XL1Blue | *recA1 endA1 gyrA96 thi-1 hsdR17 supE44 relA1 lac* [*F' proAB lacI*$^q$ *ZΔM15* Tn*10 (Tet*$^R$*)*] | [22] |
| Plasmids | | |
| pBK-miniTn7-gfp2 | pUC19-based delivery plasmid for miniTn7-*gfp2*. $Gm^R$, $Cm^R$, $Ap^R$, $mob^+$ | [23] |
| pRK600 | ColE1 RK2-$Mob^+$ RK2-$Tra^+$; helper plasmid; $Cm^R$ | [22] |
| pUX-BF13 | R6K replicon-based helper plasmid, providing the Tn7 transposition function in trans, $Ap^R mob^+$ | [22] |
| pK18-Gm-mobsacB | Suicide knockout plasmid, $Gm^R$ | [24] |
| pHerd20T | pUCP20T Plac replaced with 1.3-kb AflII-EcoRI fragment of araC-PBAD cassette, $Amp^R$ | [25] |
| pHerd20T-*siaD* | pHerd20T plasmid carrying *siaD* ORF | This study |

Table S3. The presence of c-di-GMP related genes in different *P. aeruginosa* strains. +, gene is present; -, gene is absent.

| | **PAO1** | **PA14** | **2192** | **AA2** | **AES-1R** | **AMT0023-30** | **AMT0060-3** | **AUS23** | **C3719** | **CHA** |
|---|---|---|---|---|---|---|---|---|---|---|
| *siaD* | + | + | + | + | + | + | + | + | + | + |
| *pa0290* | + | + | + | + | + | + | + | + | + | + |
| *pa0338* | + | + | + | + | + | + | + | + | + | + |
| *pa0847* | + | + | + | + | + | + | + | + | + | + |
| *roeA* | + | + | + | + | + | + | + | + | + | + |

| | | | | | | | | | | |
|---|---|---|---|---|---|---|---|---|---|---|
| *pa1851* | + | + | + | + | + | + | + | + | + | + |
| *pa2771* | + | - | + | + | + | + | + | + | + | + |
| *pa2870* | + | + | + | + | + | + | + | + | + | + |
| *pa3177* | + | + | + | + | + | + | + | + | + | + |
| *hsbD* | + | + | + | + | + | + | + | + | + | + |
| *sadC* | + | + | + | + | + | + | + | + | + | + |
| *pa4396* | + | + | + | + | + | + | + | + | + | + |
| *gcbA* | + | + | + | + | + | + | + | + | + | + |
| *pa4929* | + | + | + | + | + | + | + | + | + | + |
| *dgcH* | + | + | + | + | + | + | + | + | + | + |
| *tpbB* | + | + | + | + | + | + | + | + | + | + |
| *wspR* | + | + | + | + | + | + | + | + | + | + |
| *pa2133* | + | + | + | + | + | + | + | + | + | + |
| *pa2200* | + | + | + | + | + | + | + | + | + | + |
| *pa2572* | + | + | + | + | + | + | + | + | + | + |
| *arr* | + | - | - | - | - | - | + | - | - | - |
| *pa3825* | + | + | + | + | + | + | + | + | + | + |
| *rocR* | + | + | + | + | + | + | + | + | + | + |
| *pa4108* | + | + | + | + | + | + | + | + | + | + |
| *pa4781* | + | + | + | + | + | + | + | + | + | + |
| *pvrR* | - | + | - | - | - | - | - | - | - | - |
| *bifA* | + | + | + | + | + | + | + | + | + | + |
| *fimX* | + | + | + | + | + | + | + | + | + | + |
| *morA* | + | + | + | + | + | + | + | + | + | + |
| *mucR* | + | + | + | + | + | + | + | + | + | + |
| *pipA* | + | + | + | + | + | + | + | + | + | + |
| *rmcA* | + | + | + | + | + | + | + | + | + | + |
| *rbdA* | + | + | + | + | + | + | + | + | + | + |
| *yegE* | + | + | + | + | + | + | + | + | + | + |
| *pa1433* | + | + | + | + | + | + | + | + | + | + |
| *pa2072* | + | + | + | + | + | + | + | + | + | + |
| *pa2567* | + | + | + | + | + | + | + | + | + | + |

| | | | | | | | | | | |
|---|---|---|---|---|---|---|---|---|---|---|
| *pa3258* | + | + | + | + | + | + | + | + | + | + |
| *nbdA* | + | + | + | + | + | + | + | + | + | + |
| *dipA* | + | + | + | + | + | + | + | + | + | + |
| *proE* | + | + | + | + | + | + | + | + | + | + |
| *pa5442* | + | + | + | + | + | + | + | + | + | + |

| | **DK2** | **IST27** | **KK1** | **LES400** | **LESB58** | **NH57388A** | **NN2** | **RP1** | **TBCF10839** | **U018A** |
|---|---|---|---|---|---|---|---|---|---|---|
| *siaD* | + | + | + | + | + | + | + | + | + | + |
| *pa0290* | + | + | + | + | + | + | + | + | + | + |
| *pa0338* | + | + | + | + | + | + | + | + | + | + |
| *pa0847* | + | + | + | + | + | + | + | + | + | + |
| *roeA* | + | + | + | + | + | + | + | + | + | + |
| *pa1851* | + | + | + | + | + | + | + | + | + | + |
| *pa2771* | + | - | + | + | + | + | + | + | + | + |
| *pa2870* | + | + | + | + | + | + | + | + | + | + |
| *pa3177* | + | + | + | + | + | + | + | + | + | + |
| *hsbD* | + | + | + | + | + | + | + | + | + | + |
| *sadC* | + | + | + | + | + | + | + | + | + | + |
| *pa4396* | + | + | + | + | + | + | + | + | + | + |
| *gcbA* | + | + | + | + | + | + | + | + | + | + |
| *pa4929* | + | + | + | + | + | + | + | + | + | + |
| *dgcH* | + | + | + | + | + | + | + | + | + | + |
| *tpbB* | + | + | + | + | + | + | + | + | + | + |
| *wspR* | + | + | + | + | + | + | + | + | + | + |
| *pa2133* | + | + | + | + | + | + | + | + | + | + |
| *pa2200* | + | + | + | + | + | + | + | + | + | + |
| *pa2572* | + | + | + | + | + | + | + | + | + | + |
| *arr* | - | - | + | - | - | - | - | - | + | - |
| *pa3825* | + | + | + | + | + | + | + | + | + | + |
| *rocR* | + | + | + | + | + | + | + | + | + | + |
| *pa4108* | + | + | + | + | + | + | + | + | + | + |
| *pa4781* | + | + | + | + | + | + | + | + | + | + |

| | | | | | | | | | | |
|---|---|---|---|---|---|---|---|---|---|---|
| *pvrR* | - | - | - | - | - | - | - | - | - | - |
| *bifA* | + | + | + | + | + | + | + | + | + | + |
| *fimX* | + | + | + | + | + | + | + | + | + | + |
| *morA* | + | + | + | + | + | + | + | + | + | + |
| *mucR* | + | + | + | + | + | + | + | + | + | + |
| *pipA* | + | + | + | + | + | + | + | + | + | + |
| *rmcA* | + | + | + | + | + | + | + | + | + | + |
| *rbdA* | + | + | + | + | + | + | + | + | + | + |
| *yegE* | + | + | + | + | + | + | + | + | + | + |
| *pa1433* | + | + | + | + | + | + | + | + | + | + |
| *pa2072* | + | + | + | + | + | + | + | + | + | + |
| *pa2567* | + | + | + | + | + | + | + | + | + | + |
| *pa3258* | + | + | + | + | + | + | + | + | + | + |
| *nbdA* | + | + | + | + | + | + | + | + | + | + |
| *dipA* | + | + | + | + | + | + | + | + | + | + |
| *proE* | + | + | + | + | + | + | + | + | + | + |
| *pa5442* | + | + | + | + | + | + | + | + | + | + |

| | **BS1** | **BS2** | **BS3** | **BS5** | **BS6** | **BS9** | **BS11** | **BS14** | **OS1** | **OS2** | **OS3** | **OS4** |
|---|---|---|---|---|---|---|---|---|---|---|---|---|
| *siaD* | + | + | + | + | + | + | + | + | + | + | + | + |
| *pa0290* | + | + | + | + | + | + | + | + | + | + | + | + |
| *pa0338* | + | + | + | + | + | + | + | + | + | + | + | + |
| *pa0847* | + | + | + | + | + | + | + | + | + | + | + | + |
| *roeA* | + | + | + | + | + | + | + | + | + | + | + | + |
| *pa1851* | + | + | + | + | + | + | + | + | + | + | + | + |
| *pa2771* | + | + | + | + | + | + | + | + | + | + | + | + |
| *pa2870* | + | + | + | + | + | + | + | + | + | + | + | + |
| *pa3177* | + | + | + | + | + | + | + | + | + | + | + | + |
| *hsbD* | + | + | + | + | + | + | + | + | + | + | + | + |
| *sadC* | + | + | + | + | + | + | + | + | + | + | + | + |
| *pa4396* | + | + | + | + | + | + | + | + | + | + | + | + |
| *gcbA* | + | + | + | + | + | + | + | + | + | + | + | + |
| *pa4929* | + | + | + | + | + | + | + | + | + | + | + | + |

| | | | | | | | | | | | | |
|---|---|---|---|---|---|---|---|---|---|---|---|---|
| *dgcH* | + | + | + | + | + | + | + | + | + | + | + | + |
| *tpbB* | + | + | + | + | + | + | + | + | + | + | + | + |
| *wspR* | + | + | + | + | + | + | + | + | + | + | + | + |
| *pa2133* | + | + | + | + | + | + | + | + | + | + | + | + |
| *pa2200* | + | + | + | + | + | + | + | + | + | + | + | + |
| *pa2572* | + | + | + | + | + | + | + | + | + | + | + | + |
| *arr* | - | - | + | + | - | + | + | + | + | - | + | - |
| *pa3825* | + | + | + | + | + | + | + | + | + | + | + | + |
| *rocR* | + | + | + | + | + | + | + | + | + | + | + | + |
| *pa4108* | + | + | + | + | + | + | + | + | + | + | + | + |
| *pa4781* | + | + | + | + | + | + | + | + | + | + | + | + |
| *pvrR* | + | + | + | + | + | + | + | + | + | + | + | + |
| *bifA* | + | + | + | + | + | + | + | + | + | + | + | + |
| *fimX* | + | + | + | + | + | + | + | + | + | + | + | + |
| *morA* | + | + | + | + | + | + | + | + | + | + | + | + |
| *mucR* | + | + | + | + | + | + | + | + | + | + | + | + |
| *pipA* | + | + | + | + | + | + | + | + | + | + | + | + |
| *rmcA* | + | + | + | + | + | + | + | + | + | + | + | + |
| *rbdA* | + | + | + | + | + | + | + | + | + | + | + | + |
| *yegE* | + | + | + | + | + | + | + | + | + | + | + | + |
| *pa1433* | - | + | + | + | + | + | + | + | + | + | + | + |
| *pa2072* | + | + | + | + | + | + | + | + | + | + | + | + |
| *pa2567* | + | + | + | + | + | + | + | + | + | + | + | + |
| *pa3258* | + | + | + | + | + | + | + | + | + | + | + | + |
| *nbdA* | + | + | + | + | + | + | + | + | + | + | + | + |
| *dipA* | + | + | + | + | + | + | + | + | + | + | + | + |
| *proE* | + | + | + | + | + | + | + | + | + | + | + | + |
| *pa5442* | + | + | + | + | + | + | + | + | + | + | + | + |

Table S4. Strain cultivation time

| Strain | Aggregate culture | Planktonic culture |
|---|---|---|
| PAO1 | 6 h | 6 h |
| PA14 | 6 h | 6 h |
| 2192 | 6 h | 6 h |

| | | |
|---|---|---|
| AA2 | 6 h | 6 h |
| AMT0023-30 | 6 h | 6 h |
| AMT0060-3 | 6 h | 6 h |
| CHA | 6 h | 6 h |
| IST27N | 6 h | 6 h |
| KK1 | 6 h | 6 h |
| NN2 | 6 h | 6 h |
| OS3 | 6 h | 6 h |
| RP1 | 6 h | 6 h |
| TBCF10839 | 6 h | 6 h |
| U018A | 6 h | 6 h |
| AES-1R | 9 h | 9 h |
| AUS23 | 9 h | 9 h |
| BS1 | 9 h | 9 h |
| BS3 | 9 h | 9 h |
| DK2 | 9 h | 9 h |
| LESB58 | 9 h | 9 h |
| NH57388A | 9 h | 9 h |
| OS2 | 9 h | 9 h |
| BS2 | 12 h | 12 h |
| BS5 | 12 h | 12 h |
| BS9 | 12 h | 12 h |
| OS1 | 12 h | 12 h |
| C3719 | 18 h | 9 h |
| LES400 | 18 h | 18 h |
| OS4 | 18 h | 18 h |
| BS6 | 24 h | 24 h |
| BS11 | 24 h | 9 h |
| BS14 | 24 h | 24 h |

Table S5. MIC (µg/mL) of tobramycin and meropenem against different strains and mucoidy on PIA plates.

| | **Tobramycin** | **Meropenem** | **Mucoidy on PIA** |
|---|---|---|---|
| PAO1 | 2 | 1 | no |
| PA14 | 1 | 0.5 | no |
| 2192 | 1 | 0.5 | no |
| AA2 | 2 | 1 | no |
| AES-1R | 8 | 8 | no |
| AMT0023-30 | 1 | 0.5 | no |
| AMT0060-3 | 1 | 0.25 | no |

| | | | |
|---|---|---|---|
| AUS23 | 2 | 16 | no |
| BS1 | 1 | 1 | yes |
| BS2 | 1 | 0.125 | no |
| BS3 | 16 | 4 | yes |
| BS5 | 32 | 4 | yes |
| BS6 | 2 | 1 | yes |
| BS9 | 16 | 1 | yes |
| BS11 | 2 | 4 | yes |
| BS14 | 0.5 | 0.125 | yes |
| CHA | 2 | 0.5 | no |
| C3719 | 4 | 64 | no |
| DK2 | 32 | 32 | no |
| IST27N | 1 | 0.0625 | no |
| KK1 | 1 | 0.5 | no |
| LES400 | 8 | 8 | no |
| LESB58 | 4 | 2 | no |
| NH57388A | 2 | 0.5 | no |
| NN2 | 1 | 1 | no |
| RP1 | 1 | 1 | no |
| OS1 | 2 | 1 | yes |
| OS2 | 2 | 0.25 | yes |
| OS3 | 1 | 1 | no |
| OS4 | 0.5 | 0.125 | yes |
| TBCF10839 | 1 | 0.5 | no |
| U018A | 1 | 2 | no |

Table S6. Primers used

| Primers | Sequence 5'->3' |
|---|---|
| | Gene deletion |
| PAO1-wspR-up-F | AGCTCGGTACCCGGGATTGTTACCGTCGATATCGAGC |
| PAO1-wspR-up-R | CACCGGCTGTTCCATCAGTCGTATCAAATTGAATGCTGTTC |
| PAO1-wspR-dn-F | GAACAGCATTCAATTTGATACGACTGATGGAACAGCCGGTG |
| PAO1-wspR-dn-R | CGACGGCCAGTGCCAATCTCGATCTCGATGTTGTCGTC |
| PAO1-siaD-up-F | AGCTCGGTACCCGGGTCTCGGCCTGCTGGACATC |
| PAO1-siaD-up-R | CTGGACGCCTGAGGAGGCCAGTTGCTCCAGCGATTGCTG |
| PAO1-siaD-dn-F | CAGCAATCGCTGGAGCAACTGGCCTCCTCAGGCGTCCAG |
| PAO1-siaD-dn-R | CGACGGCCAGTGCCATTCGCCATGGAGATCATCAGC |
| PAO1-PA5442-up-F | AGCTCGGTACCCGGGATTGATCCCGAGCAACTGCTC |
| PAO1-PA5442-up-R | GAGTCGAAAGTGCTACAGCATACGAACACCAGCACCTGCTGGAG |
| PAO1-PA5442-dn-F | CTCCAGCAGGTGCTGGTGTTCGTATGCTGTAGCACTTTCGACTC |
| PAO1-PA5442-dn-R | CGACGGCCAGTGCCAAGCCTTGTCGACATTCTGC |
| | Sequencing |
| pk18-F | TGCTTCCGGCTCGTATGTTG |
| pk18-R | GCGAAAGGGGGATGTGCTG |
| pHerd20T-F | ATCGCAACTCTCTACTGTTTCT |
| pHerd20T-R | TGCAAGGCGATTAAGTTGGGT |

| | |
|---|---|
| PAO1-wspR-F | ATCGATGCAGCGGTGCAG |
| PAO1-wspR-R | AGCTGCGAAGTATACTGCACTTGC |
| PAO1-siaD-F | ACGACGAGTAGCCGACGATG |
| PAO1-siaD-R | TCGTCCTGACCCTGGAGTTG |
| PAO1-PA5442-F | ATCTGGAAGTTCTCGCTCAG |
| PAO1-PA5442-R | TTCGAGCTTCTCTTCCTCC |
| | Expression |
| SiaD-exp-F | ATGGGATCTGATAAGAATTCGTGCGGCTGGAGCGCATC |
| SiaD-exp-R | CGACGG CCA GTG CCA AGC TTT CAG CGC GCT GGA GCC GG |
| | RT-qPCR |
| qRT-siaD-F | CGCTACCAGCAGATGATG |
| qRT-siaD-R | GAGCGTTCGTTCTCCTC |
| qRT-pipA-F | ATCTCGCAATCGCCATCG |
| qRT-pipA-R | TTGCGGCGAAGAAGAACAG |
| qRT-pa0290-F | CAACGCGAATACCGCATC |
| qRT-pa0290-R | GCTTCTTGTCGGTGATGTC |
| qRT-pa0338-F | CATCCATCCCGAGGACTATC |
| qRT-pa0338-R | CGCTGATCCACAGGTAATCC |
| qRT-rmcA-F | AACCGCTTCACCTACGTC |
| qRT-rmcA-R | GTAGTGGATGGCGTACTCG |
| qRT-pa0847-F | CATGCTGTCGCCCATTAC |
| qRT-pa0847-R | TCCGACAAGAGCCAGATAC |
| qRT-rbdA-F | GGTGCTTCACGACATGAC |
| qRT-rbdA-R | TGCAGGCGATACTCGAAC |
| qRT-roeA-F | TGACCGGCCTGTTCAATC |
| qRT-roeA-R | GCGGTCGTTGATGTACTTG |
| qRT-yegE-F | GCCTGAACACCTGTTTCATC |
| qRT-yegE-R | GATCACCGAGACCAGGAAGG |
| qRT-pa1433-F | AGGCCTACCAGGACAGTCTC |
| qRT-pa1433-R | AGCCCGTTCAGGTCGTTCAG |
| qRT-pa1851-F | ATGATGGCCGAACAGGAC |
| qRT-pa1851-R | CGACATCCACCAGGATCAG |
| qRT-pa2072-F | CATCGAGCAGAGCCATTTC |
| qRT-pa2072-R | CCGGTTTCCCAGATAGACG |
| qRT-pa2133-F | GTATACCCGCGAAGCACGTC |
| qRT-pa2133-R | TCGTCCTCCTCGAGACCTTC |
| qRT-pa2200-F | TGTTGTTCGGCGTCATGCTG |
| qRT-pa2200-R | ATGGGCTGGTAGTGCACCTC |
| qRT-pa2567-F | CAATGGAACGCACGGAAC |
| qRT-pa2567-R | GGAAAGGTCGATGTCGTAGG |
| qRT-pa2572-F | CCTGCAACTGCTGGAAAG |
| qRT-pa2572-R | GTGAGCAGGATGCGAATG |
| qRT-pa2771-F | CCCTTCATCCGCTTCTACG |
| qRT-pa2771-R | ATTGAGCTGGCGTCCTTC |
| qRT-pa2870-F | CGTTCTACGCGTTGTTCTG |
| qRT-pa2870-R | TGGCGTTAAGCTCGATCAG |
| qRT-pa3177-F | GCCTGTTCTTCGAGGAAGTC |
| qRT-pa3177-R | TGTCATGGCTCAGGCGATAG |
| qRT-pa3258-F | ATTGCGACGAGAGCGTTC |
| qRT-pa3258-R | TTCGGAAAGCGAGAGGATG |
| qRT-nbdA-F | CAGATGGCTCGCTACGACAG |
| qRT-nbdA-R | TCGAGGTCGAGGAACATCAC |
| qRT-hsbD-F | ACGACTCCCGTTCCAATC |
| qRT-hsbD-R | GCTGTCCTCCAGCATGTAG |
| qRT-pa3825-F | ATCGAGTCGAGCGAAGTC |
| qRT-pa3825-R | TGGAACTTGCGCAGGTAG |
| qRT-pa4108-F | ACGTCTACGACGCGATCACC |
| qRT-pa4108-R | TTGACGAAGGCGCGGAACAC |

| | |
|---|---|
| qRT-sadC-F | CGGCATCTACCTGGTAGAG |
| qRT-sadC-R | TATGTAGCTGGCGAACAGG |
| qRT-pa4396-F | ACGGGCTCAACCTGAAG |
| qRT-pa4396-R | GCAGCAGTTCGTCGATG |
| qRT-pa4781-F | ATCTTCCGCGTATCGAGC |
| qRT-pa4781-R | GCCGATGTCATGCAACAG |
| qRT-gcbA-F | TCGATCAGCGAACTCAACC |
| qRT-gcbA-R | GTCCTTCAGTGCCAGGTAG |
| qRT-pa4929-F | CGACGAACTGCTGGAATAC |
| qRT-pa4929-R | GCACGTACCAGAAATAGGC |
| qRT-dipA-F | CAGTTGTCGCTGTCGAAG |
| qRT-dipA-R | GGCGAGTTTCTCGATGTG |
| qRT-proE-F | GACATGCAGTGGCTCAAC |
| qRT-proE-R | CGTGCTCTTCGATCAACC |
| qRT-pa5442-F | TCGCGGTGATCATGCTG |
| qRT-pa5442-R | AGGCGAAGGTGAGGAAG |
| qRT-dgcH-F | ATCGATGGCAGCCTGAAC |
| qRT-dgcH-R | CCGCAGGTATTCCTCGAAG |
| qRT-bifA-F | GGCGAGAACTTCGTCACCAC |
| qRT-bifA-R | GATCTTCGACAGCGGCTTGG |
| qRT-fimX-F | TCCTACATGGCGCTGTTC |
| qRT-fimX-R | TGGTAGCCCTTGAGGAAATC |
| qRT-morA-F | TCGTCCAGGTCAACGATTC |
| qRT-morA-R | TTGAGCTGGTTGGCTTCC |
| qRT-mucR-F | CAGCCGTACCAGATATCCC |
| qRT-mucR-R | TGGTCCTTGGCGTGATAC |
| qRT-RocR-F | CTGGTCGACAAGCTGTTC |
| qRT-RocR-R | ACCCAGTTGCGAAGGATG |
| qRT-tpbB-2-F | CTGATCGCCCGTTCCATCAG |
| qRT-tpbB-2-R | TAGACGATGGCGCTGGAGAC |
| qRT-wspR-F | GTGGCCAACCAGATCAAG |
| qRT-wspR-R | AGGACGATGATCGGGATG |
| qRT-proC-F | CAGGCCGGGCAGTTGCTGTC |
| qRT-proC-R | GGTCAGGCGCGAGGCTGTCT |
| qRT-ampC-F | AGATTCCCCTGCCTGTG |
| qRT-ampC-R | GCGGTGAAGGTCTTGCT |
| qRT-OprD-F | TCCGCAGGTAGCACTCA |
| qRT-OprD-R | AAGCCGGATTCATAGGTGG |
| qRT-recA-F | TCCGCAGGTAGCACTCAG |
| qRT-recA-R | AAGCCGGATTCATAGGTGG |
| qRT-gyrA-F | TGTGCTTTATGCCATGAGCGA |
| qRT-gyrA-R | TCCACCGAACCGAAGTTGC |
| qRT-fabD-F | GCATCCCTCGCATTCGTCT |
| qRT-fabD-R | GGCGCTCTTCAGGACCATT |
